# Confinement stress with movement restriction suppresses male courtship in *Drosophila* through dopamine-dependent neuroplasticity

**DOI:** 10.1101/2024.12.16.628796

**Authors:** Tomohito Sato, Rana Toyama, Toshihiro Kitamoto, Takaomi Sakai

## Abstract

Stress disturbs the physiological and psychological balance in animals, leading to changes in brain function. Here, we show that stress in a small space with movement restriction (SS stress) suppresses male courtship in *Drosophila* and that alterations in dopamine signaling induced by SS stress are responsible for the persistence of this suppression after the stress experience. We found that SS stress activates numerous dopamine neurons in the brain. Pharmacological and genetic analyses revealed that dopamine synthesis, release, and reception are essential for the persistence of SS-stress-induced courtship suppression. A specific subset of dopamine neurons projects to the mushroom body (MB), a brain region where various sensory inputs are integrated. We identified that SS-stress-induced dopamine release plastically depresses the activity of a subset of MB neurons, and this neuronal depression contributes to the sustained suppression of male courtship behavior following stress exposure. This novel stress model using *Drosophila* provides valuable insights into dopamine-mediated stress mechanisms, particularly those related to confined spaces.

## INTRODUCTION

Stress—whether from external or internal stimuli—can significantly disrupt an animal’s physiological and psychological balance, leading to alterations in brain function and overall physiological state. Experiencing such stress conditions induces plastic changes in the brain that may persist after the stress experience, affecting both physiology and behavior (*1*). In animals, including humans, both acute and chronic stressors have been shown to affect various behavioral outputs, including the wake–sleep cycle (*2*), emotional behavior (*3, 4*), escape responses (*5, 6*), locomotion (*7, 8*), and feeding patterns (*9, 10*).

Under stress conditions, neurotransmitter release and signaling are altered, leading to diverse physiological and behavioral stress responses (*11*). Dopamine plays a significant role in these processes. In humans, acute stress has been shown to trigger dopamine release in the prefrontal cortex (PFC), which is associated with impaired working memory performance in males. This suggests that increased dopamine secretion in the PFC impairs PFC function during a stress experience (*12*). On the other hand, long-term exposure to psychosocial adversity has been linked to reduced dopaminergic activity in the striatum (*13*). These findings indicate that stress can both enhance and suppress dopaminergic function, depending on its nature and duration. In rodent models, chronic stress (e.g., social defeat, isolation, restraint, or exposure to aversive odors or environments) leads to long-lasting deleterious effects on dopamine function in several brain regions and on dopamine-related behaviors, mirroring findings in humans (*12*). Interestingly, stress-induced modifications of dopaminergic function are not limited to vertebrates but are also observed in invertebrates. For instance, mechanical stress results in a four-fold increase in dopamine levels in the hemolymph of oysters (*14*). In the fruitfly *Drosophila*, dopamine levels rise following short-term (30–60 min) exposure to heat stress (*15, 16*), and mechanical stress increases the enzymatic activity of tyrosine hydroxylase (TH), the rate-limiting enzyme for dopamine synthesis, but starvation stress decreases TH activity (*17*). Furthermore, dopamine levels are reduced in ants subjected to isolation stress and in bees exposed to mechanical stress (*18, 19*). These observations highlight the widespread effects of stress on dopaminergic function across a wide range of species, from vertebrates to invertebrates.

Sexual behavior is an instinctive and essential activity of many sexually dimorphic animal species, particularly those with a well-developed nervous system, as it plays a critical role in species conservation. Similar to other intrinsic behaviors, sexual behavior in mammals can be affected by stress. For example, patients with post-traumatic stress disorder, which can result from traumatic experiences such as combat or accidents, often exhibit sexual dysfunction (*20*). Similarly, male rats show reduced sexual motivation following exposure to stressors such as foot shock and predator odors (*21, 22*). Despite these observations, the molecular and cellular mechanisms by which stress diminishes sexual motivation remain poorly understood. *Drosophila*, a model organism with well-established neurogenetic tools and behavioral assay, has become a valuable system for studying how stress affects brain function and behavior (*7, 23*). *Drosophila* also has a rich history in sexual behavior research (*24–26*), with significant progress in identifying the neural mechanisms that regulate sexual behavior (*27–29*). Notably, dopamine neurons and dopamine release have been found to be critical for controlling male mating drive and sexual motivation (*30, 31*). Thus, neurogenetic studies using *Drosophila* offer a promising approach to uncovering the mechanism underlying stress-induced reduction in sexual motivation and exploring the role of dopamine in stress responses.

In this study, we established a novel experimental paradigm to induce stress in *Drosophila* by confining a male fly in a very tight space using a small acrylic chamber [hereafter referred to as small space (SS) stress]. Under the SS stress condition, males are unable to walk freely but can freely move their legs and turn their bodies within the chamber. In this regard, this setup differs from the restraint stress methods commonly used in rodent studies. Using this stress paradigm, we found that male courtship activity toward virgin females was significantly suppressed following exposure to SS stress. Furthermore, we demonstrated that dopamine signaling plays a role in this stress-induced suppression of male courtship activity.

## RESULTS

### Courtship suppression persists for at least 1 h after the experience of 1 h SS stress

To expose flies to SS stress, a sexually mature virgin male (4–6 d old) of the wild-type strain Canton-S (CS) was placed in a tiny chamber (3 mm diameter × 2 mm depth) for 10, 30, and 60 min (Fig. 1A, SS chamber). As a control, a virgin male was placed in a larger chamber (15 mm diameter × 3 mm depth) where it could walk freely (Fig. 1A, standard chamber). Hereafter, males exposed to SS stress are called “stressed males”, whereas control males are referred to as “naive males”. After the stress experience, tests were performed using naïve and stressed males. In the tests, male courtship behavior was observed immediately (0 h) following the SS stress experience (Fig. 1B), and then male courtship activity in both naive and stressed males was quantified using the courtship index (CI), defined as the percentage of time spent performing courtship behaviors over a 10 min period (Fig. 1B, CI). The performance index (PI) was subsequently calculated as an indicator of the degree of courtship suppression (Fig. IB, PI). Males subjected to 30 or 60 min of SS stress exhibited a significantly lower courtship activity than naive males (Fig. 1C, CI), indicating that SS stress of these durations induces male courtship suppression. However, no significant difference in the CI was detected between naive and stressed males after 10 min of SS stress (Fig. 1C, CI), suggesting that 10 min of SS stress is insufficient to induce courtship suppression. Furthermore, the effect of stress-induced courtship suppression weakened progressively as the duration of SS stress decreased (Fig. 1C, PI). On the basis of these findings, we primarily used 1 h SS stress in subsequent experiments.

**Fig. 1.**
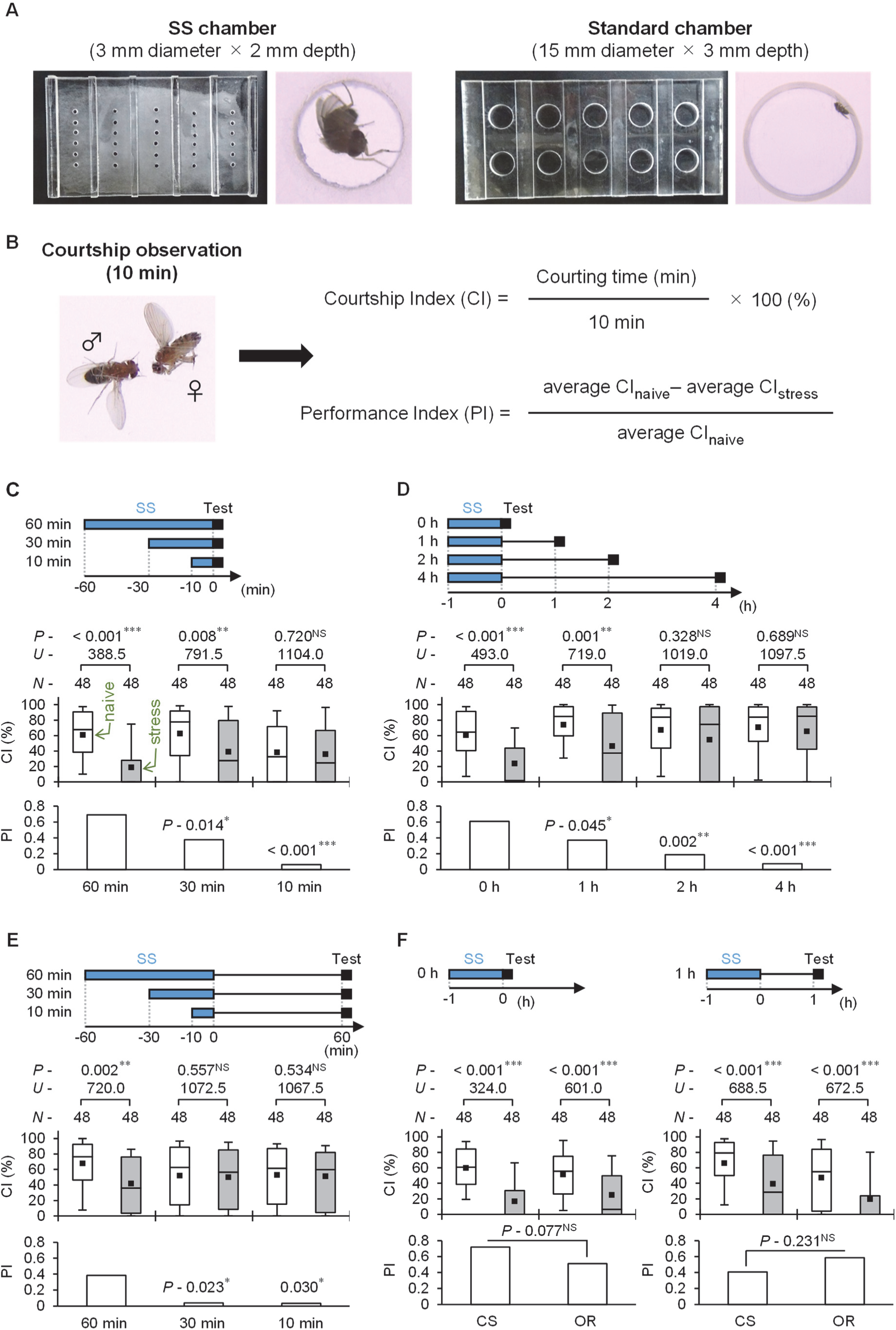
Male courtship suppression after the experience of SS stress. (**A**) Acrylic chambers were used in the SS stress assay. Single males were introduced into the SS chamber to expose the flies to SS stress (left). Naive males introduced into the standard chamber were used as a control (right). (**B**) Male courtship behavior was observed after SS stress in each experiment. Male courtship behavior was observed as previously described (*65*). Single intact virgin male and freeze-killed virgin female couples were transferred into observation chambers (15 mm diameter, 3 mm depth), and courtship behaviors were videotaped for 10 min. The formulas for determining the courtship index (CI) and performance index (PI) are shown on the right side of the diagram. For more details, see Materials and Methods. (**C**) Courtship activity was measured immediately after 60, 30, and 10 min SS stress. (**D**) Courtship activity was measured immediately, 1 h, 2 h, and 4 h after 1 h SS stress. (**E**) Courtship activity was measured 60 min after 60, 30, and 10 min SS stress. (**C** to **E**) Wild-type (CS) males were used in the experiments. (**F**) Courtship activity in two wild-type strains (CS and OR) was measured immediately (0 h) and 1 h after 1 h SS stress. (**C** to **F**) In each graph, white boxes indicate naive males and gray boxes indicate stressed males. Box plots for a set of CI data show the 10th, 25th, 75th, and 90th centiles. In the box plots, black squares indicate the mean, and the lines are drawn at the median. For statistical comparisons, the Mann–Whitney *U*-test was used for CI, and a bootstrapping-based randomization test with an R-script (*67*) was used for PI. *N*, sample size; ***, *P* < 0.001; **, *P* < 0.01; *, *P* < 0.05; NS, not significant.

Next, male courtship activity was measured immediately (0 h), 1, 2, and 4 h after 1 h SS stress (Fig. 1D). Stressed males showed significantly lower courtship activity than naive males immediately and 1 h after 1 h SS stress (Fig. 1D, CI). However, no significant difference in the CI was detected between naive and stressed males 2 and 4 h after SS stress (Fig. 1D, CI). These findings indicate that the effect of stress-induced courtship suppression diminishes over time after the SS stress experience (Fig. 1D, PI). Additionally, we confirmed no courtship suppression 1 h after 10 or 30 min of SS stress (Fig. 1E). These findings indicate that 1 h SS stress is necessary to induce male courtship suppression, which persists for at least 1 h after the stress experience. To assess whether the fly genetic background affects the courtship suppression induced by 1 h SS stress, we conducted the same experiments using another wild-type strain, Oregon-R (OR). OR males also showed significant courtship suppression immediately and 1 h after 1 h SS stress (Fig. 1F). Thus, regardless of the genetic background of wild-type strains, 1 h SS stress consistently induces male courtship suppression.

In *Drosophila*, repeated exposure to vibration stress has been shown to induce a low motivational state, affecting various behaviors such as climbing, spontaneous locomotor activity, appetite, and sexual desire (*7*). To determine whether SS stress affects behaviors beyond male courtship, we measured spontaneous locomotor activity and appetite following 1 h SS stress. Locomotor activity in stressed males was significantly reduced immediately after 1 h SS stress (Fig. S1A). However, no significant difference was detected between naive and stressed males 1 h after the SS stress exposure (Fig. S1A), indicating that courtship suppression observed 1 h after 1 h SS stress is not simply due to general sluggishness. Next, we assessed feeding behavior as an indicator of appetite following 1 hr of SS stress. Using an automatic feeding monitoring system, FlyPAD (*32*), we measured the number of sips and sip duration in individual males. Males were fasted for 23 h, and feeding behavior was monitored for 1 h after 1 h SS stress. No significant differences were detected in either the number of sips or sip duration between naive and stressed males after the experience of 1 h SS stress (Fig. S1, B), indicating that the stress has minimal impact on male appetite.

We further examined whether longer stress experiences extend the duration of subsequent courtship suppression. Male courtship activity was measured on day 1 (d 1) and day 5 (d 5) following 7 h and 24 h SS stress. On the day after the 7 h and 24 h SS stress, male courtship activity was significantly reduced (Fig. S2A), despite no observable reduction in locomotor activity (Fig. S2B). Moreover, this courtship suppression persisted for at least 5 days (Fig. S2A). These findings suggest that the duration of courtship suppression is affected by the length of the stress experience.

### Dopamine plays a role in the persistence of courtship suppression following SS stress

To investigate the involvement of dopamine in stress-induced courtship suppression, we used 3-iodo-L-tyrosine (3IY)-fed males in the experiments. 3IY inhibits tyrosine hydroxylase (TH), the enzyme that catalyzes the conversion of L-tyrosine to a dopamine precursor, thereby disrupting dopamine synthesis (*33*). In *Drosophila*, 3IY feeding reduces dopamine levels in both larvae and adults (*34, 35*). Virgin males (3 d old) were fed 3IY (0.1 or 1 mg/ml) for 2 days before the experiments. Feeding 3IY did not affect courtship suppression immediately after 1 h SS stress (Fig. 2A, left). However, males fed with 3IY did not show courtship suppression 1 h after 1 h SS stress (Fig. 2A, right). These findings suggest that dopamine synthesis is primarily responsible for maintaining courtship suppression after the experience of SS stress.

**Fig. 2.**
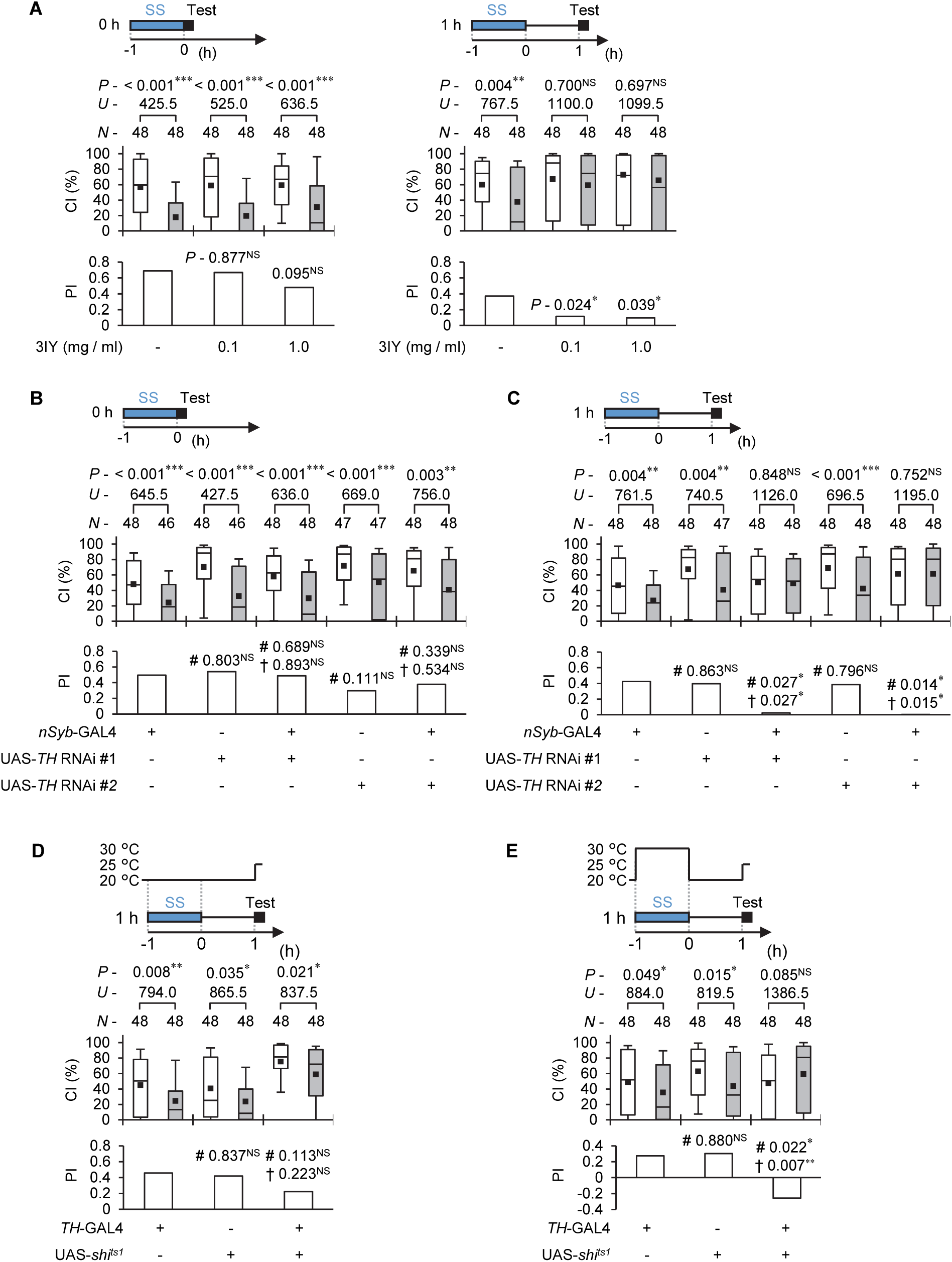
Dopamine is involved in the persistence of courtship suppression after SS stress experience. (**A**) Wild-type (CS) males with or without 3IY-feeding were used in the experiments. Courtship activity was measured immediately (0 h) and 1 h after 1 h SS stress. 3IY-feeding was conducted for 2 d before the experiments. (**B** and **C**) Male courtship activity was measured 1 h after 1 h SS stress using TH knockdown flies. *TH*-RNAi #1 or *TH*-RNAi #2 was driven by *nSyb*-GAL4. Courtship activity was measured immediately (**B**) and 1 h (**C**) after 1 h SS stress. (**D** and **E**) *shi^ts1^* was driven by *TH*-GAL4. Virgin males were collected and kept at 20 °C until the experiments. (**D**) After 1 h SS stress at PT (20 °C) followed by 1 h maintenance of stressed flies at 20 °C, male courtship activity was measured at 25 °C. (**E**) After 1 h SS stress at RT (30 °C) followed by 1 h maintenance of stressed flies at 20 °C, male courtship activity was measured at 25 °C. (**A** to **E**) In each graph, white boxes indicate naive males, and gray boxes indicate stressed males. Box plots for a set of CI data show the 10th, 25th, 75th, and 90th centiles. In the box plots, black squares indicate the mean, and the lines are drawn at the median. For statistical comparisons, the Mann–Whitney *U*-test was used for CI, and a bootstrapping-based randomization test with an R-script (*67*) was used for PI. # and† indicate *P*-values against GAL4 and UAS controls, respectively. *N*, sample size; *** *P* < 0.001; **; *P* < 0.01; *, *P* < 0.05; NS, not significant.

We next investigated whether the pan-neural knockdown of TH affects courtship suppression following 1 h SS stress. For this, we used a pan-neural GAL4 line, *nSyb*-GAL4, and two UAS-*TH* RNAi lines [UAS-*TH* RNAi #1 (TRiP HMS05881) and UAS-*TH* RNAi #2 (TRiP HMC06137)]. Immunostaining adult brain neurons with an anti-TH antibody revealed that TH expression in the adult brain was almost completely absent in TH knockdown flies (Fig. S3). Similar to the inhibition of dopamine synthesis by 3IY, TH knockdown flies (*nSyb*-GAL4/UAS-*TH* RNAi) did not exhibit courtship suppression 1 h after 1 h SS stress (Fig. 2B). However, they did display courtship suppression immediately after 1 h SS stress (Fig. 2C). These findings suggest that dopamine synthesis in neurons is essential for maintaining courtship suppression following the experience of SS stress, but is not required for the initial induction of courtship suppression immediately after SS stress.

### SS stress modifies the activity of dopamine neurons in the brain

Does SS stress increase or decrease the activity of dopamine neurons in the fly brain? To answer this question, we measured the activity of dopamine neurons during and after SS stress exposure. In *Drosophila*, genetic tools for measuring lasting changes in neuronal intracellular calcium levels are often used to estimate neuronal excitability (*36–38*). To examine whether dopamine neurons are activated by SS stress, we used a calcium-modulated photoactivatable ratiometric integrator (CaMPARI2), which changes its fluorescence from green to red only in the presence of high Ca^2+^ levels and user-supplied UV light (*38, 39*). In *Drosophila*, the adult brain contains approximately 300 dopamine neurons (*40, 41*). We focused on nine neuronal clusters composed of multiple neurons (PAM, PAL, PPM1, PPM2, PPM3, PPL1, PPL2ab, PPL2c, and T1). To express CaMPARI2 in dopamine neurons, we used *TH*-GAL4/UAS-*CaMPARI2* males. The fluorescence intensities of CaMPARI2 red and green signals were measured under UV light in each cluster following 1 h SS stress. In the PPL1, PPL2c, and PPM1 clusters, the red-to-green ratio in each dopamine neuron in stressed males was significantly higher than that in naive males (Fig. 3A), likely reflecting intracellular Ca^2+^ levels increased by the stress condition. In contrast, the red-to-green ratio in the PPL2ab neurons was lower in stressed males than in naive males (Fig. 3B), suggesting that SS stress inhibits the activity of PPL2ab neurons. No significant differences in the red-to-green ratio were observed in the PAL, PPM2, PPM3, and T1 clusters between stressed and naive males (Fig. 3C), indicating that these clusters of dopamine neurons are minimally affected by 1 h SS stress.

**Fig. 3.**
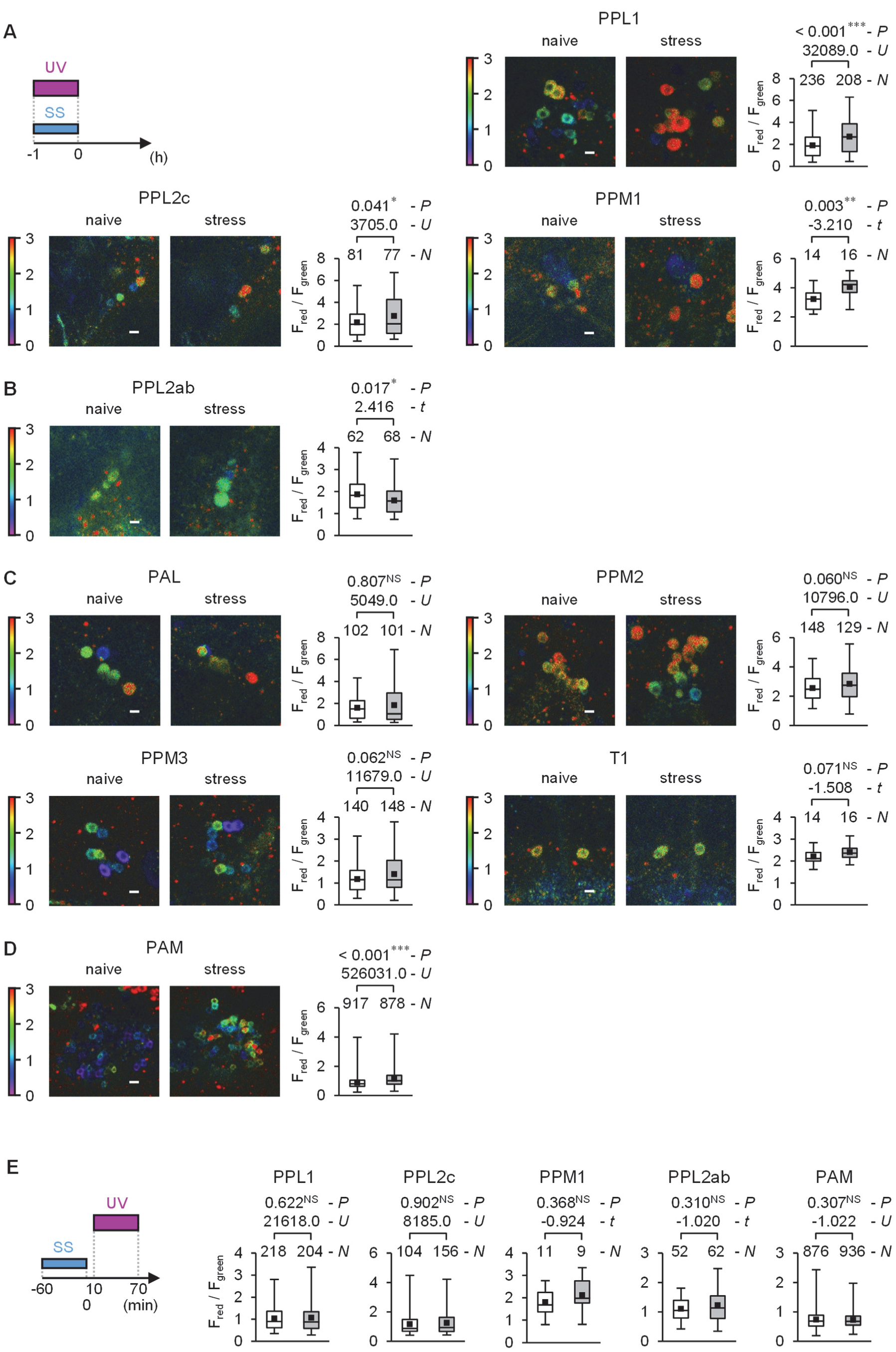
SS stress modifies the activity of dopamine neurons in the brain. (**A to C**) CaMPARI2 was used to compare dopamine neuron activity between control and stressed males. 395 nm UV light (15.0 ± 0.2 mW/cm^2^) was irradiated during 1 h SS stress. CaMPARI2 was driven by *TH*-GAL4. (**A**) Box plots show the quantified ratio of green to red signals in PPL1, PPL2c, and PPM1. (**B**) Box plots show the quantified ratio of green to red signals in PPL2ab. (**C**) Box plots show the quantified ratio of green to red signals in PAL, PPM2, PPM3, and T1. (**D**) Box plots show the quantified ratio of green to red signals in PAM. In this experiment, CaMPARI2 was driven by *R58E02*. The UV light intensity was 20.0 ± 0.2 mW/cm^2^. (**E**) UV light irradiation (20.0 ± 0.2 mW/cm^2^ for PPL1, PPL2c, PPM1, and PPL2ab; 15.0 ± 0.2 mW/cm^2^ for PAM) was conducted for 10–70 min after 1 h SS stress. (**A** to **E**) Heat maps indicate the red/green fluorescence ratio in the representative confocal image of each dopamine cluster. Scale bars indicate 5 µm. In each graph, white boxes indicate naive males, and gray boxes indicate stressed males. Box plots for red/green fluorescence ratio data show the 0th, 25th, 75th, and 100th centiles. In the box plots, black squares indicate the mean, and the lines are drawn at the median. Round ROIs were set on each soma. Six to ten brains were used. The Mann–Whitney *U*-test, Student’s t-test or Welch’s t-test was used for statistical comparisons. *N*, sample size (cell numbers); ***, *P* < 0.001; **, *P* < 0.01; *, *P* < 0.05; NS, not significant.

There are approximately 100 dopamine neurons in the PAM cluster, but in *TH*-GAL4 lines, only 13 PAM neurons per hemisphere express GAL4 (*40*). To address this limitation, we used the *R58E02* line, which drives GAL4 expression in approximately 80% of PAM dopamine neurons (*42*). In stressed males, the red-to-green ratio in PAM dopamine neurons was significantly higher than that in naive males (Fig. 3D). In summary, results of our experiments using CaMPARI2 demonstrate that 1 h SS stress increases the activity of dopamine neurons in at least four neuronal clusters in the brain (PAM, PPL1, PPL2c, and PPM1) and reduces the activity in a small subset of dopamine neurons (PPL2ab).

Next, we investigated whether intracellular Ca^2+^ levels in dopamine neurons remain elevated after 1 h SS stress. Flies were exposed to UV light for 1 h, beginning 10 min after the SS stress experience and continuing until 70 min post-stress. Unlike during SS stress, intracellular Ca^2+^ levels in dopamine neurons in stressed males were comparable to those in naive males (Fig. 3E). These findings suggest that in the PAM, PPL1, PPL2c, and PPM1 clusters, dopamine neuron activity is elevated during SS stress but returns to baseline levels after the SS stress experience.

### Neurotransmission from dopamine neurons during exposure to SS stress contributes to the persistence of courtship suppression after the stress experience

CaMPARI2 imaging revealed that dopamine neurons in the PAM, PPL1, PPL2c, and PPM1 clusters are activated during 1 h SS stress (Fig. 3), suggesting that dopamine release during this time might underlie the persistence of male courtship suppression after the stress. To test this hypothesis, we conducted experiments to inhibit neurotransmission from dopamine neurons during SS stress exposure. For these experiments, we used the temperature-sensitive Dynamin mutation *shibire^ts1^* (*shi^ts1^*), expressed in dopamine neurons via the *TH*-GAL4 driver and UAS-*shi^ts1^*lines. *TH*-GAL4 can be used as a GAL4 driver for dopamine neurons because the GAL4 expression pattern in *TH*-GAL4 is similar, if not identical, to that of endogenous TH (*40*). Shi^ts1^ can inhibit neurotransmission in a temperature-dependent manner (*43*). To disrupt neurotransmission during SS stress, 1 h SS stress was performed at a restrictive temperature (RT) of 30 °C. Afterward, flies were kept at a permissive temperature (PT) of 20 °C for 1 h, and then male courtship activity was measured at a PT of 25 °C. When 1 h SS stress was performed at PT (20 °C), *TH*-GAL4/UAS-*shi^ts1^*males showed courtship suppression, similarly to control flies (*TH*-GAL4/+ and UAS-*shi^ts1^*/+) (Fig. 2D). However, when 1 h SS stress was performed at RT (30 °C), *TH*-GAL4/UAS-*shi^ts1^* males did not show courtship suppression. Under this condition, control males still exhibited courtship suppression (Fig. 2E, RT). These findings indicate that dopamine release during SS stress is critical for the persistence of male courtship suppression following stress exposure.

### Three types of dopamine receptor are responsible for the persistence of courtship suppression after stress experience

*Drosophila* has four dopamine receptors, all of which are G protein-coupled receptors: Dop1R1, Dop1R2, Dop2R, and DopEcR (*37, 44*). Dop1R1 and Dop1R2 are D1-like receptors that activate the cAMP signaling pathway (*37*), whereas Dop2R is a D2-like receptor believed to inhibit the cAMP pathway (*37*). Additionally, flies have a noncanonical dopamine receptor, DopEcR, which is activated by both dopamine and ecdysone (*37*). To determine which dopamine receptors are required for male courtship suppression after SS stress exposure, we used knockout (KO) GAL4 lines for each dopamine receptor (*45*). These KO GAL4 lines are null mutants, as one or more exons of each dopamine receptor gene are replaced by GAL4. Courtship suppression immediately after 1 h SS stress was detected in all four KO GAL4 lines (Fig. 4A), indicating that the absence of any single dopamine receptor does not affect the immediate response. However, when examining the persistence of courtship suppression, we found that homozygotes or hemizygous males of three KO GAL4 males (*Dop1R1^KOGAL4^*, *Dop1R2^KOGAL4^*, and *Dop2R^KOGAL4^*) did not exhibit courtship suppression after 1 h SS stress. In contrast, males lacking DopEcR (*DopEcR^KOGAL4^*) continued to show courtship suppression (Fig. 4B). These findings demonstrate that Dop1R1, Dop1R2, and Dop2R are necessary for maintaining courtship suppression after the experience of SS stress.

**Fig. 4.**
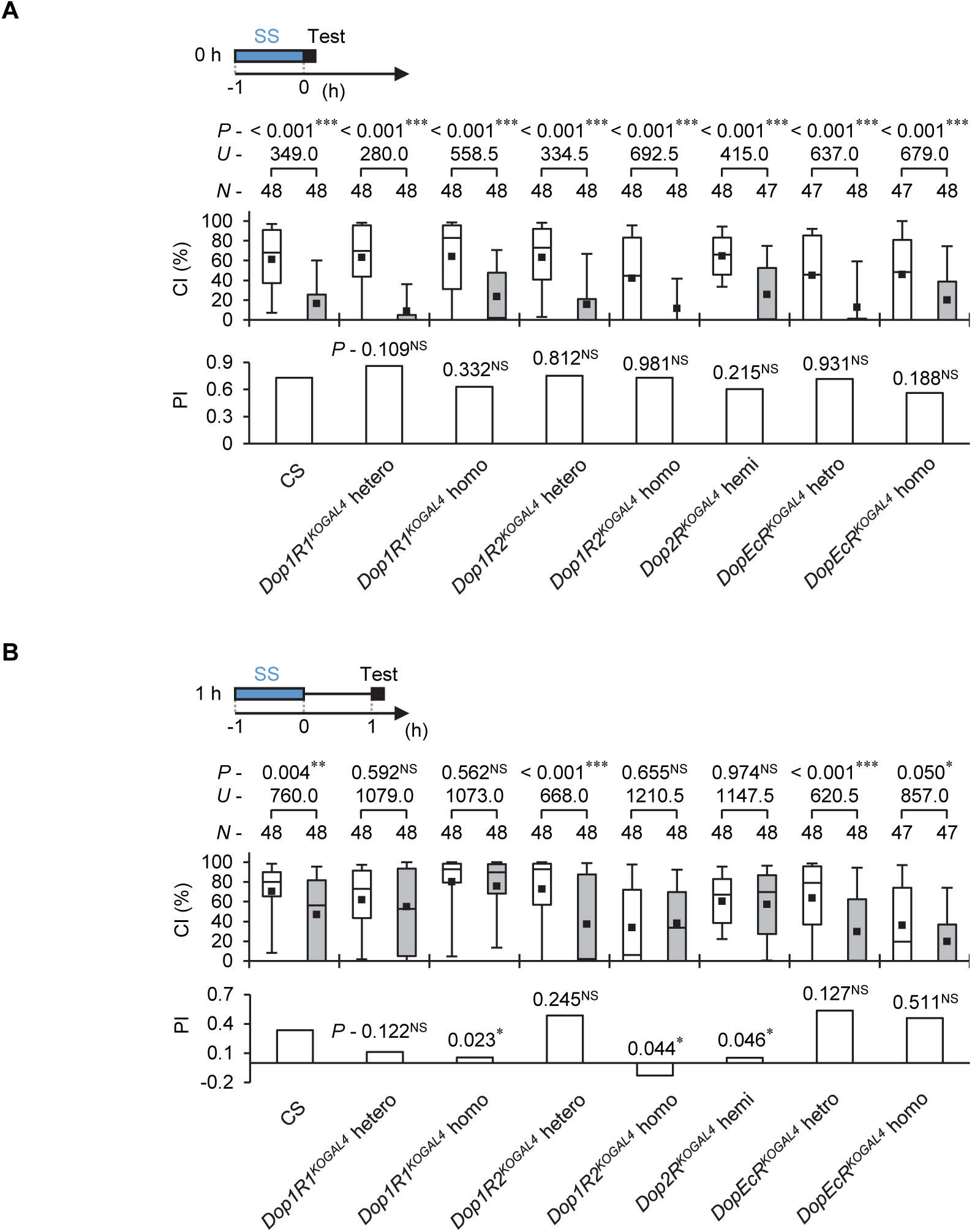
Dopamine receptors are responsible for the persistence of courtship suppression after SS stress experience. (**A**) Courtship activity was measured immediately after 1 h SS stress. (**B**) Courtship activity was measured 1 h after 1 h SS stress. (**A** and **B**) GAL4 knockout lines of four dopamine receptor genes were used in the experiments (*Dop1R1^KOGAL4^*, *Dop1R2^KOGAL4^*, *Dop2R^KOGAL4^*, and *DopEcR^KOGAL4^*). In each graph, white boxes indicate naive males and gray boxes indicate stressed males. Box plots for a set of CI data show the 10th, 25th, 75th, and 90th centiles. In the box plots, black squares indicate the mean, and the lines are drawn at the median. In statistical analyses, the Mann–Whitney *U*-test was used for CI, and a bootstrapping-based randomization test with an R-script (*67*) was used for PI. For PI, each genotype was compared with wild-type (CS). *N*, sample size; ***, *P* < 0.001; **, *P* < 0.01; *, *P* < 0.05; NS, not significant.

### Dopamine receptors in mushroom body (MB) neurons are involved in the persistence of courtship suppression after stress experience

Neurons in the PAM, PPL1, and PPL2ab clusters terminate and innervate MB neurons, which are required for various *Drosophila* behaviors (*46*). Furthermore, Dop1R1, Dop1R2, and Dop2R are expressed in MB neurons (*45, 47*). To examine whether dopamine receptors in MB neurons contribute to stress-induced courtship suppression, we performed RNA interference (RNAi) experiments targeting these receptors. We used the UAS-*Dop1R1* RNAi, UAS-*Dop1R2* RNAi, and UAS-*Dop2R* RNAi lines in combination with the pan-MB GAL4 line *R13F02*. The TRiP RNAi line UAS-*Dop1R2* RNAi, constructed using the VALIUM20 vector, is effective without enforced expression of *Dicer2* (https://bdsc.indiana.edu/stocks/rnai/rnai_all.html); thus, this line was used independently of UAS-*Dicer2*. Conversely, the other UAS-RNAi lines were combined with UAS-*Dicer2* to enhance RNAi efficiency. The knockdown effectiveness of all three UAS-RNAi lines was confirmed by qRT-PCR using the pan-neural *nSyb*-GAL4 line (Fig. S4). Behavioral analysis revealed that males with *DopR1*or *Dop2R* knockdown did not show courtship suppression 1 h after 1 h SS stress, whereas males with *Dop1R2* knockdown retained this suppression (Fig. 5A). These results indicate that Dop1R1 and Dop2R in MB neurons are required to maintain courtship suppression following SS stress.

**Fig. 5.**
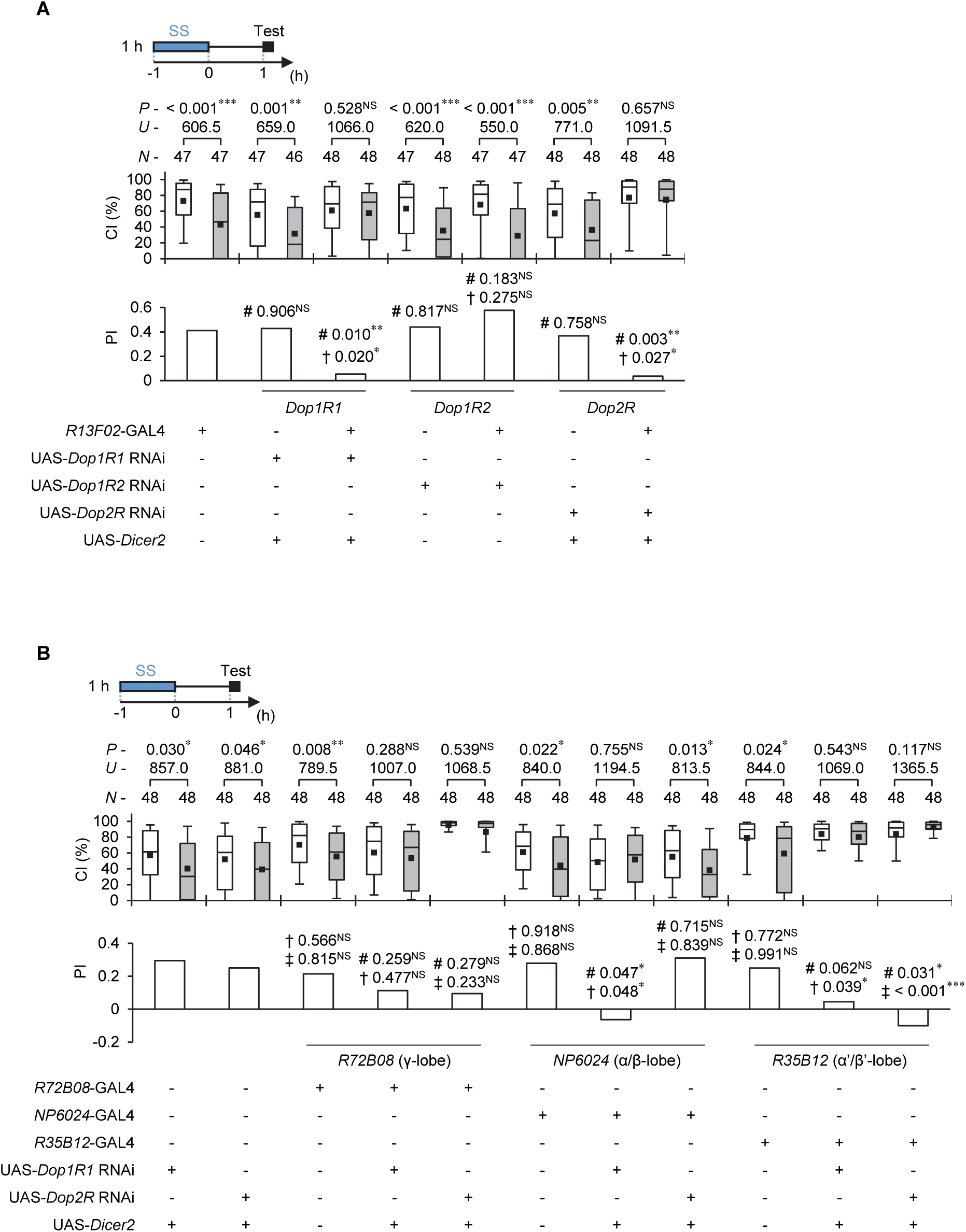
Dopamine receptors in the mushroom body are involved in the persistence of courtship suppression after SS stress experience. (**A** and **B**) Male courtship activity was measured 1 h after 1 h SS stress. Males with *Dop1R1*, *Dop1R2*, or *Dop2R* knockdown were used. In each graph, white boxes indicate naive males and gray boxes indicate stressed males. Box plots for a set of CI data show the 10th, 25th, 75th, and 90th centiles. In the box plots, black squares indicate the mean, and the lines are drawn at the median. As statistical comparisons, the Mann–Whitney *U*-test was used for CI, and a bootstrapping-based randomization test with an R-script (*67*) was used for PI. Number signs (#) indicate *P*-values against the GAL4 control (**A** and **B**),† indicate *P*-values against UAS controls (**A**), and †and‡indicate *P*-values against UAS controls of the *Dop1R1* RNAi and *Dop2R* RNAi, respectively (**B**). *N*, sample size; ***, *P* < 0.001; **, *P* < 0.01; *, *P* < 0.05; NS, not significant. (A) *Dop1R1* RNAi, *Dop1R2* RNAi, and *Dop2R* RNAi were driven by a pan-MB GAL4 line, R13F02. (B) *Dop1R1* RNAi and *Dop2R* RNAi were driven by MB γ-GAL4 (*R72B08*), MB α/β -GAL4 (*NP6024*), and MB α’/β’-GAL4 (*R35B12*).

To identify the specific MB neurons critical for SS-stress-induced courtship suppression, we used three GAL4 lines to target distinct MB neuronal subsets: *NP6024* for MB α/β neurons, *R72B02* for MB γ neurons, and *R35B12* for MB α’/β’ neurons. Male courtship activity was assessed 1 h after 1 h SS stress. In males with *Dop1R1* or *Dop2R* knockdown in MB γ neurons, there were no significant differences in the CI between naive and stressed males (Fig. 5B, γ), although their PI was not significantly different from that of control males. These findings suggest that Dop1R1 and Dop2R in MB γ neurons partially contribute to courtship suppression. In contrast, *Dop1R1* knockdown in MB α/β neurons abolished SS-stress-induced courtship suppression (Fig. 5B, α/β). However, *Dop2R* knockdown in MB α/β neurons had no effect (Fig. 5B, α/β), indicating that Dop1R1 is the primary receptor involved in this neuronal subset for maintaining courtship suppression 1 h after 1 h SS stress exposure. In MB α’/β’ neurons, the knockdown of either *Dop1R1* or *Dop2R* prevented male courtship suppression (Fig. 5B, α’/β’). These findings indicate that MB αβ, γ, and α’β’ neurons each play a role in SS-stress-induced courtship suppression, but the contribution of specific dopamine receptors varies among these neuronal types.

### Neurotransmission in PAM and PPL1 is involved in the persistence of courtship suppression after stress experience

Since PAM, PPL1, and PPL2ab dopamine neurons terminate and innervate MB neurons (*46*), we next examined whether inhibiting neurotransmitter release from these clusters during SS stress exposure could prevent the persistence of courtship suppression after SS stress experience. We used two GAL4 lines (*R58E02* for PAM and *NP5945* for PPL2ab) and a split-GAL4 line (*MB504B* for PPL1). In *R58E02*/UAS-*shi^ts1^*and *MB504B*/UAS-*shi^ts1^* males, no courtship suppression was observed when SS stress was applied at RT (30 °C) (Fig. 6A). In contrast, suppression was present when the stress was conducted at PT (25 °C) (Fig. 6B). These results suggest that dopamine release from PAM and PPL1 neurons is critical for maintaining courtship suppression for at least 1 h after 1 h SS stress. Unlike *R58E02* and *MB504B* lines, *NP5945*/UAS-*shi^ts1^* males showed courtship suppression at both PT and RT (Figs. 6A and 6B), indicating that blocking dopamine release from PPL2ab has little effect on SS stress-induced courtship suppression.

**Fig. 6.**
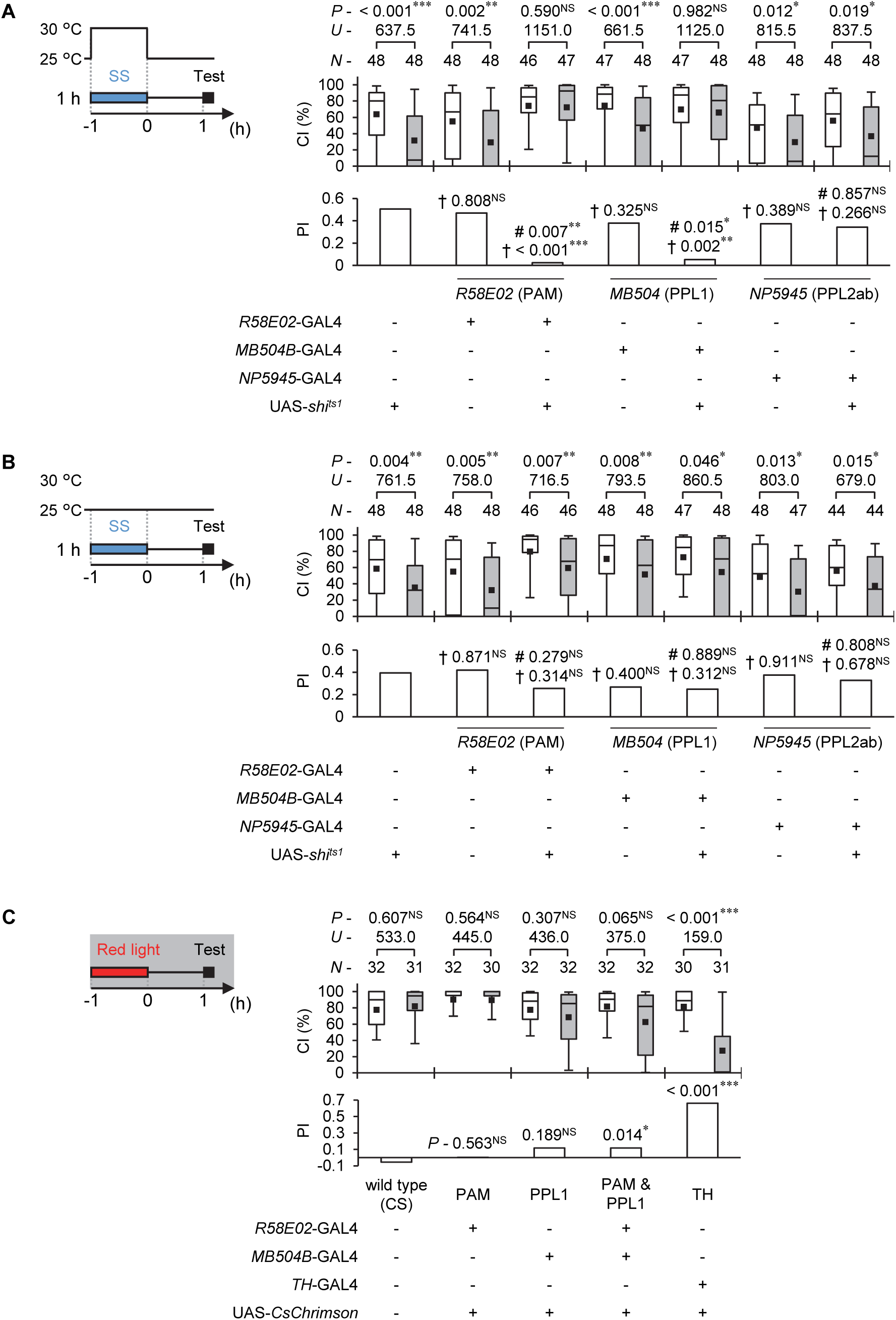
Disruption of neurotransmission in PAM and PPL1 prevents the persistence of courtship suppression after SS stress experience. To induce *shi^ts1^* expression in PAM, PPL1, and PPL2ab, three GAL4 lines were used (*R58E02* for PAM, *MB504B* for PPL1, and *NP5945* for PPL2ab). (**A**) SS stress was applied at 30 °C for 1 h followed by the maintenance of stressed flies at 25 °C for 1 h, and then the courtship observation was conducted at 25 °C. (**B**) SS stress was applied at 25 °C for 1 h followed by the maintenance of stressed flies at 25 °C for 1 h, and then the courtship observation was conducted at 25 °C. (**A** and **B**) In each graph, white boxes indicate naive males and gray boxes indicate stressed males. Box plots for a set of CI data show the 10th, 25th, 75th, and 90th centiles. In the box plots, black squares indicate the mean, and the lines are drawn at the median. For statistical comparisons, the Mann–Whitney *U*-test was used for CI, and a bootstrapping-based randomization test with an R-script (*67*) was used for PI. Number signs (#) and daggers (†) indicate *P*-values against GAL4 and UAS controls, respectively. (**C**) Optogenetic activation of dopamine neurons induces courtship suppression. Wild-type (CS) males and males expressing *CsChrimson* expression in a subset of dopamine neurons were used in the experiments. Three GAL4 lines (*R58E02* for PAM, *MB504B* for PPL1, and *TH*-GAL4 for almost dopamine neurons) were used to induce *CsChrimson* in dopamine neurons. Male courtship activity was measured 1 h after 1 h red light irradiation (630 nm, 1000 ± 50 µW/cm^2^). In each graph, white boxes indicate naive males and gray boxes indicate ATR-fed males. Box plots for a set of CI data show the 10th, 25th, 75th, and 90th centiles. In the box plots, black squares indicate the mean, and the lines are drawn at the median. For statistical comparisons, the Mann– Whitney *U*-test was used for CI. For PI, a bootstrapping-based randomization test with an R-script was used. (**A** to **C**) *N*, sample size; ***, *P* < 0.001; **, *P* < 0.01; *, *P* < 0.05; NS, not significant.

### Activation of dopamine neurons triggers the persistence of courtship suppression

As shown in Fig. 3, many dopamine neurons in at least four clusters are activated following 1 h SS stress. To investigate whether artificial activation of dopamine neurons can induce courtship suppression, we used the optogenetic tool CsChrimson, a red-shifted channelrhodopsin (ChR) with strong light sensitivity in the 510–640 nm range (*48, 49*). CsChrimson expression activates target neurons in a red-light-dependent manner. The efficiency of ChR activation depends significantly on the amount of supplemented all-*trans* retinal (ATR) (*48*). To ensure ATR feeding itself does not affect male courtship activity, we compared males orally administered ATR (ATR+) with those not fed ATR (ATR-) in semidarkness (< 2.0 µW/cm^2^). We confirmed that ATR feeding does not affect male courtship activity (Fig. S5). Next, we examined whether activation of either PAM or PPL1 neurons induces courtship suppression. For this, we used *R58E02*/UAS-*CsChrimson* and *MB504B*/UAS-*CsChrimson* males. After 1 h of red light irradiation, we compared courtship activity between ATR+ and ATR-males. Similarly to wild-type (CS) males, ATR+ *R58E02*/UAS-*CsChrimson* and *MB504B*/UAS-*CsChrimson* males did not exhibit courtship suppression 1 h after 1 h red light irradiation, as their courtship activities were comparable to those of ATR-males (Fig. 6C). We then assessed whether simultaneous activation of PAM and PPL1 neurons could induce courtship suppression. However, even with concurrent activation, ATR+ males did not show courtship suppression 1 h after 1 h red light irradiation (Fig. 6C, CI). Finally, to test whether activating a broader population of dopamine neurons induces courtship suppression, we used *TH*-GAL4/UAS-*CsChrimson* males. In this case, ATR+ males showed reduced courtship activity compared with ATR-males 1 h after 1 h red light irradiation (Fig. 6C). These findings indicate that the activation of multiple dopamine neuron clusters, not only PAM and PPL1, is indispensable for inducing courtship suppression lasting at least 1 h.

### SS stress induces plastic changes in the activity of MB γ neurons, and these changes contribute to the persistence of courtship suppression following stress experience

Our three experiments—3IY intake, TH knockdown, and dopamine receptor knockdown— demonstrated that dopamine signaling plays a critical role in sustaining male courtship suppression after SS stress (Figs. 2 and 5). Notably, during SS stress exposure, the activation of dopamine neurons in four clusters was prominent (Fig. 3). Additionally, experiments using *shi^s1^* revealed that dopamine release during SS stress exposure is essential for the persistence of male courtship suppression (Fig. 2). From on these findings, it seems logical that dopamine release is promoted by the activation of dopamine neurons during stress exposure and that dopamine levels should return to the baseline after the stress experience (Fig. S6A). However, signaling via dopamine receptors appears necessary to sustain male courtship suppression after the stress experience, rather than during the experience. This raises an important question: why does male courtship suppression persist after SS stress even though dopamine release is no longer activated? We propose the following hypothesis: Active dopamine release during SS stress exposure may induce plastic alterations in the function of the dopamine-receptor-expressing neurons in the brain. These plastic changes likely persist after the stress experience, leading to continued male courtship suppression. To test this hypothesis, we conducted experiments to determine whether the activity of MB neurons, which express dopamine receptors, undergoes plastic changes after SS stress exposure and whether these changes depend on dopamine signaling.

We measured intracellular Ca^2+^ levels in MB neurons to investigate whether the excitability of these neurons undergoes plastic changes as a result of SS stress experience. To compare Ca^2+^ levels in the MB α/β, γ, and α’/β’ lobes in males with and without stress exposure, we performed two types of experiment using UAS-*CaMPARI2* in combination with three MB GAL4 lines (*NP6024* for MB α/β neurons, *R72B02* for MB γ neurons, and *R35B12* for MB α’/β’ neurons). First, to determine whether MB activity increases during SS stress, flies were irradiated with UV light during a 1 h SS stress session, and the red-to-green ratio of CaMPARI2 was measured immediately afterward. In all MB lobes except for the α’ lobe, stressed males exhibited a significantly higher red-to-green ratio than naive males (Fig. 7A-C). This indicates that intracellular Ca^2+^ levels in the MB α/β, γ, and β’ lobes increase under stress conditions. However, similar enhancements of CaMPARI2 responses were observed in 3IY-fed males, suggesting that the rise in Ca^2+^ levels during SS stress is independent of dopamine signaling (Fig. 7D). Next, to evaluate Ca^2+^ levels after the SS stress experience, flies were irradiated with UV light for 1 h, starting 10 min after the SS stress session. In the MB α/β, γ, and β’lobes, there were no significant differences in Ca^2+^ levels between naïve and stressed males (Fig. 7E). However, in the MB γ lobe, stressed males showed a reduction in Ca^2+^ levels compared with naïve males within the first hour after SS stress (Fig. 7F). The reduced Ca^2+^ levels returned to the baseline 2 h after the SS stress experience (Fig. 7G). Notably, in 3IY-fed flies, no reduction in Ca^2+^ levels in the MB γ lobe was observed 1 h after SS stress (Fig. 7F), indicating that this reduction depends on dopamine signaling. In summary, most MB lobes exhibit increased activity during SS stress in a dopamine-independent manner. However, the MB γ lobe uniquely shows a dopamine-dependent reduction in activity that persists for at least 1 h after SS stress and subsequently returns to the baseline (Fig. S6B).

**Fig. 7.**
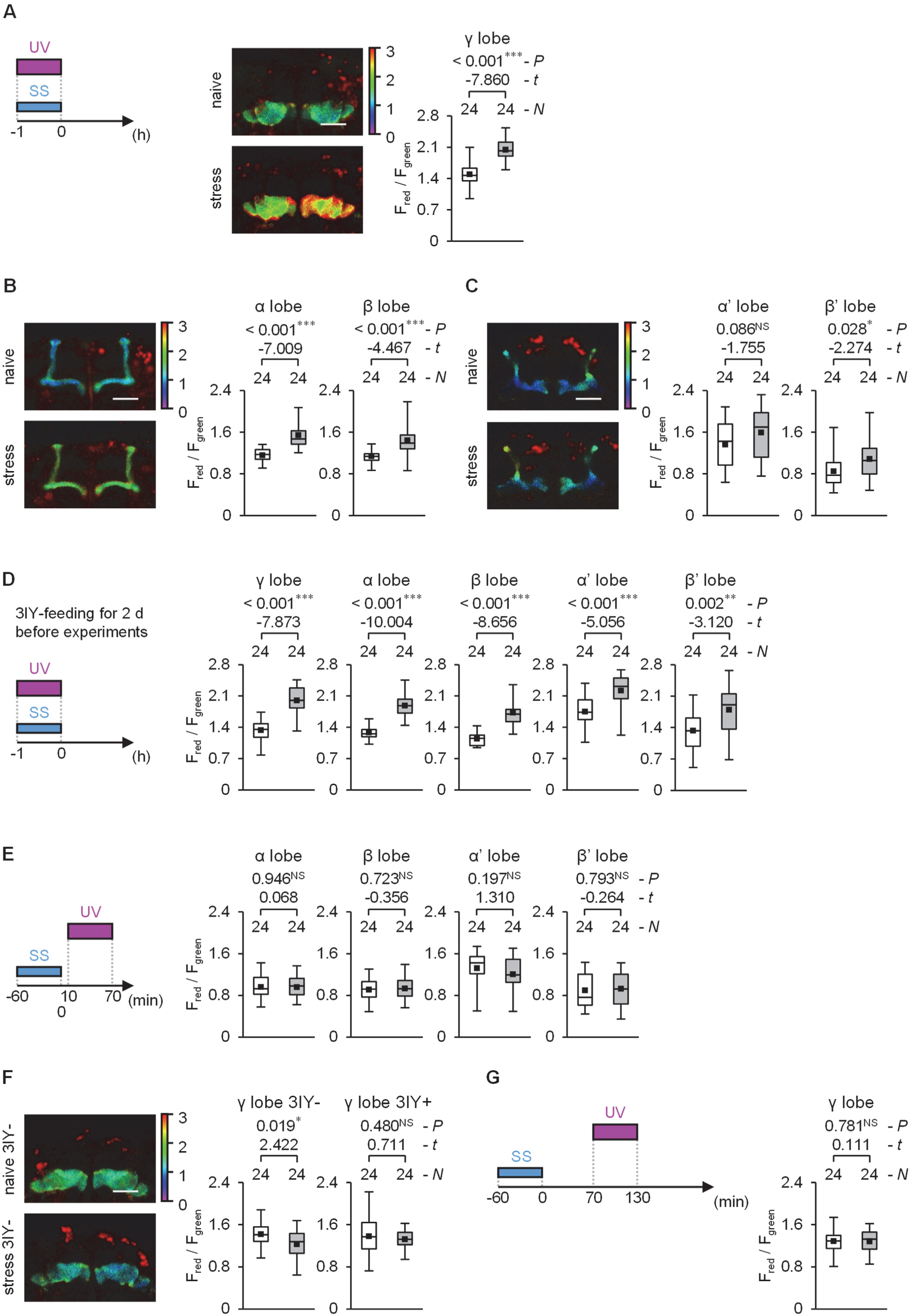
SS stress modifies the activity of MB neurons. (**A** to **F**) Box plots show the quantified ratio of green to red signals in MB γ (**A**), α/β (**B**), and α’/β’ (**C**) lobes. Three GAL4lines (*R72B02*, *NP6024*, and *R35B12*) were used to drive *CaMPARI2* in MB γ, α/β, and α’/β’ neurons. UV light (395 nm, 10.0 ± 0.2 mW/cm^2^) was used for the photoconversion of CaMPARI2. (**A** to **C**) Males that were not fed 3IY were used. UV light was irradiated during 1 h SS stress. Heat maps indicate the red/green fluorescence ratio in the representative confocal image of each MB lobe. Scale bars indicate 50 µm. (**D**) Males were used in the experiments. Males were fed 3IY for 2 d before the experiments. UV light was irradiated during 1 h SS stress. (**E**) Males were not fed 3IY were used. UV light was applied from 10 min to 70 min after 1 h SS stress. (**F**) 3IY-fed males and non-3IY-fed males were used in the experiments. Males were fed 3IY for 2 d before the experiments. (**G**) UV light was irradiated from 70 to 130 min after 1 h SS stress. (**A** to **G**) In each graph, white boxes indicate naive males and gray boxes indicate stressed males. Box plots for red/green fluorescence ratio data show the 0th, 25th, 75th, and 100th centiles. In the box plots, black squares indicate the mean, and the lines are drawn at the median. Polygonal ROIs were set on each entire lobe. Sample sizes indicate lobe numbers. Twelve brains were observed in all experiments. For statistical comparisons, the Student’s *t*-test or Welch’s *t*-test was used on the basis of homoscedasticity. *N*, sample size; ***, *P* < 0.001; **, *P* < 0.01; *, *P* < 0.05; NS, not significant.

Does the reduction in Ca^2+^ levels in MB γ neurons after experiencing SS stress contribute to male courtship suppression? If so, activating MB γ neurons with CsChrimson after 1 h SS stress should prevent male courtship suppression. To test this hypothesis, we irradiated *R72B02*/UAS-*CsChrimson* males with red light for 1 h, starting immediately after the 1 h SS stress experience and continuing until immediately before the courtship test begins. Courtship activity was then measured in semidarkness (Fig. 8, red light). In these experiments, we used flies that had been orally administered ATR (ATR+) as well as those that had not (ATR-). Among ATR+ males exposed to red light, there was no significant difference in CI between stressed and nonstressed males, indicating that male courtship suppression was blocked (Fig. 8, red light, ATR+). However, in ATR-males, male courtship suppression was still detected (Fig. 8, red light, ATR+). Additionally, we performed a control experiment in which ATR+ males were not exposed to red light. These males were kept in a dark box for 1 h immediately after the SS stress experience and then tested for courtship activity in semidarkness (Fig. 8, dark). Under these experimental conditions, the CI of the stressed ATR+ males was significantly lower than that of the nonstressed ATR+ males, clearly indicating male courtship suppression (Fig. 8, dark). These findings suggest that neuronal activation by CsChrimson compensates for the stress-induced depression of MB γ neurons, thereby preventing male courtship suppression following SS stress. Thus, the stress-induced depression of MB γ neurons appears to play a key role in the persistence of male courtship suppression after stress experience.

**Fig. 8.**
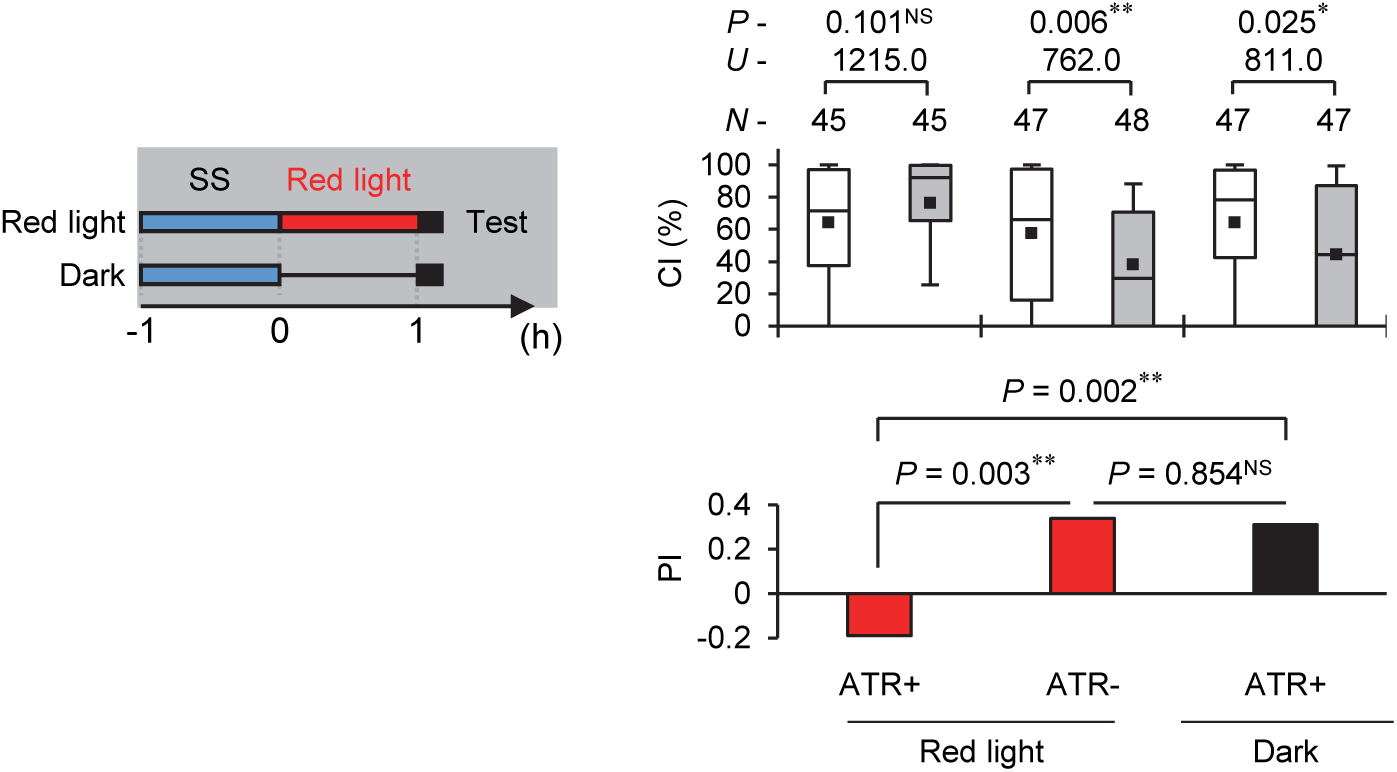
Optogenetic activation in MB γ neurons after the SS stress experience prevents male courtship suppression. Males expressing CsChrimson in MB γ-neurons were used in the experiments. *R72B02* was used as the MB γ-neuron GAL4 line. 10.0 ± 0.5 µW/cm^2^ red light was irradiated on ATR+ males and ATR-males for 1 h from immediately after 1 h SS stress to before the test initiation. As a control for red light irradiation, we used ATR+ males kept in the dark for 1 h after the SS stress experience. In the test, male courtship activity was observed in semidarkness (< 2.0 µW/cm^2^). In each graph, white boxes indicate naive males and gray boxes indicate stressed males. Box plots for a set of CI data show the 10th, 25th, 75th, and 90th centiles. In the box plots, black squares indicate the mean, and the lines are drawn at the median. In statistical comparisons, the Mann–Whitney *U*-test was used for CI. For PI, a bootstrapping-based randomization test with an R-script (*67*) was used to compare between conditions. *N*, sample size; **, *P* < 0.01; *, *P* < 0.05; NS, not significant.

## DISCUSSION

Although stress experiences often lead to sexual dysfunctions (e.g., difficulties of sexual desire, orgasm, and satisfaction) in mammals (*20, 22, 50*), the molecular and cellular mechanisms underlying these effects remain unclear. In this study, we demonstrated for the first time that *Drosophila* males show courtship suppression after experiencing SS stress. A 1 h SS stress exposure suppressed male courtship activity for at least 1 h (Fig. 1), whereas longer SS stress exposures of 7 h and 24 h induced courtship suppression lasting at least 5 d (Fig. S2). These findings suggest that the duration of courtship suppression after a stressful experience is directly affected by the length of the stress experience. In *Drosophila*, exposure to stressors such as starvation or heat shock does not significantly affect male courtship activity (*51, 52*). However, Rise et al. reported that flies exposed to vibration stress (300 Hz vibration for 10 h per day) exhibit reduced motivation for behaviors related to sexual desire, general locomotion, and appetite (*7*). Unlike their finding, we observed that spontaneous locomotor activity and appetite remained unaffected 1 h after 1 h SS stress (Fig. 1 and Fig. S1). These differences suggest that distinct stress types or variations in the duration of the stress experience differentially impact brain function and behavior.

We found that the temporal inhibition of TH by 3IY, neuron-specific TH knockdown, and temporal disruption of neurotransmission in TH-positive cells all had no effect on male courtship suppression immediately after SS stress but prevented courtship suppression at 1 h after the SS stress experience. These findings indicate that dopamine synthesis and release during SS stress are essential for the persistence of courtship suppression following the stress experience (Fig. 2). Similarly, knockout mutations of three dopamine receptors prevented courtship suppression 1 h after SS stress but had no effect immediately after the SS stress experience (Fig. 4). These results suggest that dopamine signaling plays a key role in maintaining courtship suppression for at least 1 h after SS stress. However, the mechanisms underlying the immediate reduction in courtship activity following SS stress remain unknown. Identifying the neurotransmitters that act as the initial triggers for reduced courtship activity during SS stress would provide valuable insights into these mechanisms.

In dopamine neurons in PAM, PPL1, PPL2c, and PPM1 clusters, intracellular Ca^2+^ levels were high in stressed flies (Fig. 3). In contrast, intracellular Ca^2+^ levels were low in the PPL2ab cluster under the same conditions (Fig. 3). In *Drosophila*, previous studies have shown that different types of stress can either increase or decrease dopamine levels, suggesting that dopamine levels are modulated in a stress-type-specific manner (*15–17*). Our findings indicate that even when exposed to the same stress, individual dopamine neurons in the brain may respond differently. Therefore, measuring cell-specific responses to stress is crucial for understanding the mechanisms underlying stress-induced modifications in dopamine signaling and their impact on behaviors.

PAM, PPL1, and PPL2ab clusters project to MB neurons (*40*). Intracellular Ca^2+^ levels in PAM and PPL1 clusters increased during the SS stress experience (Fig. 3). However, these levels returned to the baseline after the SS stress exposure (Fig. 3 and Fig. S6), suggesting that SS stress acutely activates these two dopamine neuron clusters. Disruption of neurotransmission from these neurons revealed that dopamine release from PAM and PPL1 clusters is necessary for courtship suppression following the SS stress experience (Fig. 7). In contrast to PAM and PPL1, neuronal activity in the PPL2ab cluster is acutely suppressed during the SS stress experience (Fig. 3). However, disruption of neurotransmission from the PPL2ab cluster during the SS stress experience did not affect male courtship suppression (Fig. 7), suggesting that PPL2ab activity contributes minimally, if at all, to the persistence of male courtship suppression after SS stress. Further experiments using males with knockdown of dopamine receptors revealed that Dop1R1 in MB α/β, γ, and α’/β’ neurons and Dop2R in MB α’/β’ and γ neurons play a predominant role in maintaining courtship suppression after SS stress (Fig. 5). Given that dopamine release was activated during 1 h SS stress exposure and returned to the baseline afterward (Fig. S6), Dop1R1 and Dop2R in the MB neurons were likely activated during SS stress exposure. These findings collectively suggest that dopamine transmission from PAM and PPL1 clusters to MB lobes during stress exposure is crucial for the persistence of male courtship suppression after the SS stress experience.

Although neurotransmitter release from PAM or PPL1 clusters is necessary to maintain courtship suppression for at least 1 h after SS stress (Fig. 7), photoactivation of PAM or PPL1, or both clusters for 1 h failed to sustain courtship suppression following optogenetic activation (Fig 8). These findings suggest that dopamine release from PAM and PPL1 alone is insufficient to maintain courtship suppression. However, photoactivation of multiple dopamine neuron clusters using the *TH*-GAL4 driver was sufficient to maintain courtship suppression (Fig. 8). This indicates that sustaining courtship suppression after SS stress requires dopamine release not only from neurons projecting to MB neurons but also from other dopamine neurons that do not target the MB neurons. Although all three dopamine receptors (Dop1R1, Dop1R2, and Dop2R) are required for maintaining SS-stress-induced courtship suppression (Fig. 4), Dop1R1 and Dop2R are predominantly involved in MB neurons. This suggests that Dop1R2 may be expressed in non-MB neurons that play a role in SS-stress-induced courtship suppression. Unlike PAM and PPL1 neurons, PPL2c and PPM1 neurons are unlikely to innervate MB neurons (*40*). Since these two clusters exhibited significantly increased intracellular Ca^2+^ levels in response to SS stress, it will be intriguing to identify the non-MB interneurons activated by SS stress that are targeted by PPL2c and PPM1 neurons. Investigating the molecular and cellular mechanisms through which these neurons suppress male courtship activity could provide deeper insights into the pathways underlying stress-induced behavioral changes. Figure 9 provides a schematic of our overall findings in this study and a proposed model based on these findings.

**Fig. 9.**
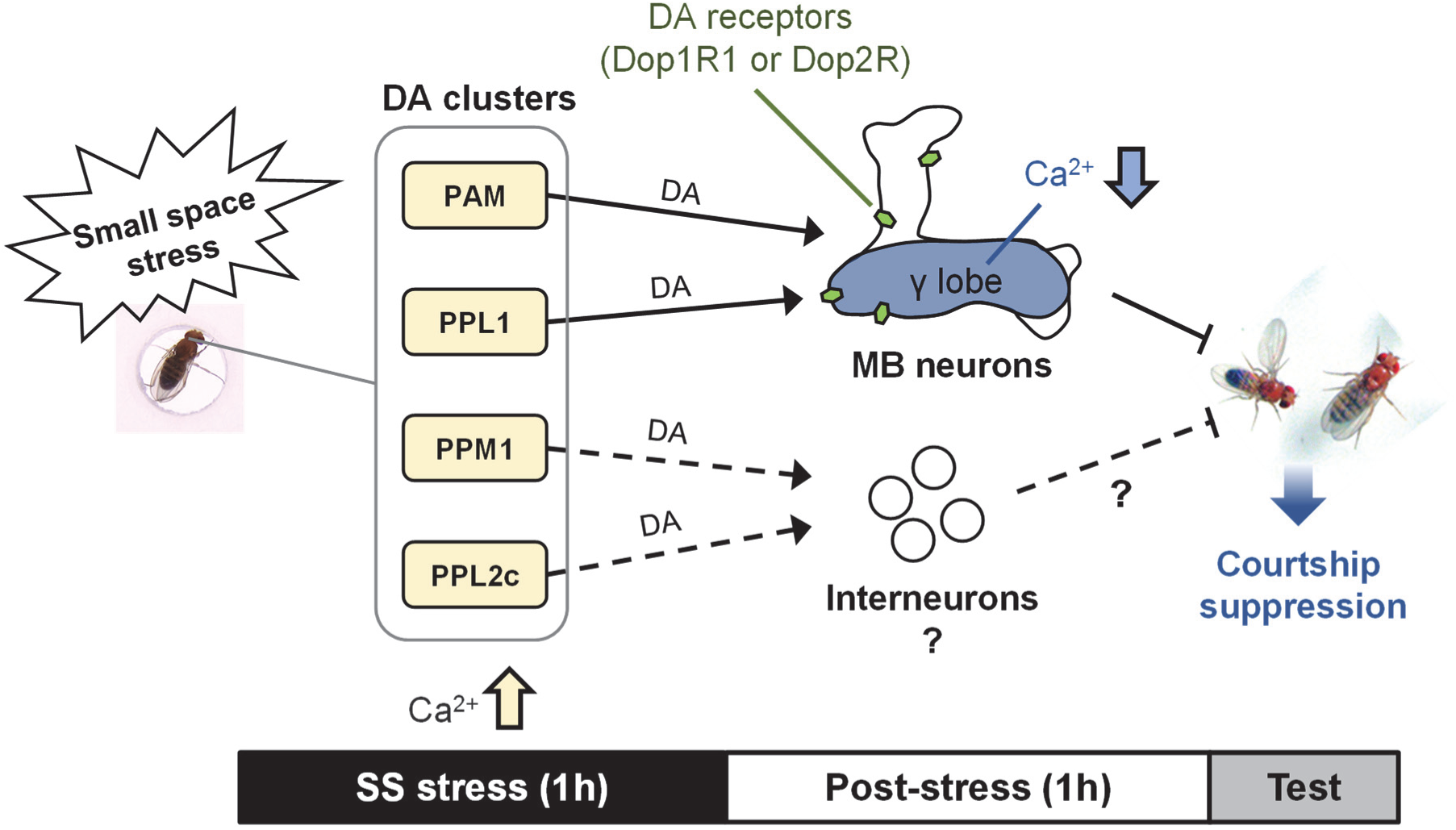
Possible model of dopaminergic functions in stress-dependent courtship suppression. The activities of PAM, PPL1, PPM1, and PPL2c are promoted during the exposure to SS stress. Dop1R1 and Dop2R in MB are necessary for stress-dependent courtship suppression. Dopamine release from PAM and PPL1 to MB induces neuronal depression in the MB γ lobe. Dopamine signals from PPM1 and PPL2c should be transmitted to non-MB neurons. Thus, PAM/PPL1/MB signaling and PPM1/PPL2c/non-MB signaling may contribute to the persistence of male courtship suppression after SS stress exposure. DA, dopamine.

In this study, we confirmed that MB neuronal activity is modified both during and after the experience of SS stress. Although Ca^2+^ levels increased in most MB lobes, in the axonal regions of MB neurons, during SS stress in a dopamine-independent manner (Fig. 7), they returned to the baseline after the SS stress experience (Fig. 7). In contrast, Ca^2+^ levels in the MB γ lobes decreased for approximately 1 h after SS stress in a dopamine-dependent manner (Fig. 7). This reduction in Ca^2+^ levels in the MB γ lobes contributes to the persistent suppression of male courtship after stress (Fig. 8). The calcium dynamics observed in MB γ-neurons following SS stress are unique and not observed in other MB neurons. This dopamine-dependent phenotype indicates that SS stress plastically alters the physiological properties of MB γ neurons via dopamine signaling. These plastic changes are responsible for the persistent suppression of male courtship after stress. However, the precise mechanism by which dopamine signaling suppresses the activity of MB γ neurons following SS stress remains unclear. In isolated cultured *Drosophila* brains, repetitive stimulation of the antennal lobe has been shown to elevate cAMP and Ca^2+^ levels in MB neurons, subsequently leading to suppressed MB neuronal activity (*53*). Similarly, elevated cAMP and Ca^2+^ levels in MB neurons during SS stress may alter their physiological properties. This hypothesis is supported by the fact that Dop1R1 activation is crucial for cAMP production in MB neurons (*54–56*), whereas Dop2R activation is considered to increase intracellular Ca^2+^ levels in MB γ neurons (*55*). Future studies should aim to elucidate how Dop1R1 and Dop2R function in MB neurons to suppress MB γ neuron activity following SS stress. Understanding these mechanisms will provide further insights into the role of dopamine signaling in stress-induced behavioral changes.

In rodents, acute and chronic immobilization stress caused by confinement in a small tube affects various behaviors, including anxiety and depression-like behaviors (*57–59*). During immobilization, rodents experience anxiety and fear owing to their inability to walk freely and the difficulty of limb movement. As a result, this stress assay with an animal model is widely used for studying human mental disorders. In mice, acute immobilization stress suppresses escape-related behaviors during the tail suspension test, and the D3-type dopamine receptor is involved in this stress response (*57*). Additionally, acute and chronic immobilization stresses increase *TH* mRNA expression levels in the locus ceruleus of rats (*60*). These findings suggest that immobilization stress alters dopaminergic functions in rodents. In the SS stress model used in this study, flies are unable to walk freely but can move their legs and body axis. Although the effects of the experimental environment differ between immobilization stress in rodents and SS stress in flies, both stress paradigms share the common features of restricting free walking and altering dopamine signaling in the brain. This suggests that changes in dopamine signaling in the brain induced by restricted walking may represent a shared phenomenon across vertebrates and invertebrates. Moreover, restraint stress has been shown to induce sexual dysfunction in male rats, including reduced sexual motivation and erectile dysfunction (*61, 62*). These findings suggest that dopamine signaling may also play a role in restraint-stress-induced sexual dysfunction in male rats.

In this study, we established a novel paradigm for studying stress in *Drosophila*. What are the sources of stress under SS conditions? One likely source is psychological stress caused by the inability to satisfy the innate desire for free walk. It is well known that restricted movement due to a small cage or enclosure size can act as a significant stressor for animals (*63*). However, the physiological mechanisms by which such stress affects brain function remain poorly understood. Similarly, solitary confinement has been shown to have detrimental effects on mental health in humans (*64*), but the underlying mechanisms are not well understood. Our findings using SS stress in *Drosophila* provide valuable new insights into how confinement and restricted movement can induce stress. This research has the potential to deepen our understanding of stress mechanisms across species, including humans, particularly in contexts involving limited physical freedom.

## MATERIALS AND METHODS

### Fly stocks

All flies were raised on glucose–yeast–cornmeal medium in 12:12 LD cycles at 25.0 ± 0.5 °C (60 ± 20% relative humidity). In the behavioral analysis and Ca^2+^ imaging experiments, virgin males and females were collected within 8 h after eclosion without anesthesia. Each virgin male was isolated until experiments except for feeding assay.

The fly stocks used for this study were as follows: Canton-S (CS), Oregon-R (OR), *nSyb*-GAL4 [51635, Bloomington Drosophila Stock Center (BDSC)], *TH*-GAL4 (8848, BDSC), *R58E02* (41347, BDSC), *MB504B* (gifted from Dr. Nagano, Tokyo Metropolitan Institute of Medical Science), *NP5945* [105062, Drosophila Genomics and Genetic Resources (DGGR)], *R13F02* (48571, BDSC), *NP6024* (105080, DGGR), *R35B12* (49822, BDSC), *R72B08* (46669, BDSC), *Dop1R1^KOGal4^* (84714, BDSC), *Dop1R2^KOGal4^* (84715, BDSC), *Dop2R^KOGal4^* (84716, BDSC), *DopEcR^KOGal4^* (84717, BDSC), UAS-*TH* RNAi #1 (76069, BDSC), UAS-*TH* RNAi #2(65875, BDSC), UAS-*Dop1R1* RNAi (93708, BDSC), UAS-*Dop1R2* RNAi (51423, BDSC), UAS-*Dop2R* RNAi (93711, BDSC), UAS-*Dicer2* (24650, BDSC), UAS-*mCD8::GFP*, UAS-*CaMPARI2* (78317, BDSC), UAS-*shi^ts1^*(*43*), and UAS-*CsChrimson* (55135 and 55136, BDSC). All lines were backcrossed to *white^1118^* flies with the CS background for at least five generations except for UAS-*mCD8::GFP*, UAS-*TH* RNAi #1, UAS-*TH* RNAi #2, UAS-*Dop1R2* RNAi, *DopEcR^KOGal4^*, and Oregon-R. In the backcrossing of *DopEcR^KOGal4^*, GAL4 insertion was confirmed in each generation by PCR. Primer sequences used in the backcrossing are as follows: Forward, 5′-AAAGAAAAACCGAAGTGCGCC-3′; Reverse, 5′-GGTCCGTTTTCAGGAAGGGC-3′.

The TRiP RNAi lines generated using the VALIUM20 vector used in this study (UAS-*TH* RNAi #1 UAS-*TH* RNAi #2, and UAS-*Dop1R2* RNAi) are expected to be effective even without being combined with UAS-*Dicer2* (https://bdsc.indiana.edu/stocks/rnai/rnai_all.html), so they were used without being combined with UAS-*Dicer2*. The other UAS-RNAi lines were used in combination with UAS-*Dicer2*.

### Small space stress assay

3- to 6-d-old virgin males were used in the experiments. All experimental procedures were conducted at 25 ± 0.5 °C (60 ± 20% relative humidity) except for the experiments using UAS*-shi^ts1^*. For cold anesthesia, males were collected into a chilled glass vial on ice within 2–3 minutes. Subsequently, each male was transferred into each well of the standard chambers (15 mm diameter, 3 mm depth, Fig. 1A, right pictures) or SS chambers (3 mm diameter, 2 mm depth, Fig. 1A, left pictures). For 7 h and 24 h SS stress, males were fed on fly foods in the chambers to avoid desiccation and hunger. Except for the case of immediate tests, males were kept in small rearing vials (10 mm diameter, 75 mm height) until the tests after the SS manipulation.

### Behavioral analyses (male courtship activity, spontaneous locomotor activity, and feeding activity)

Tests of male courtship activity were carried out as previously described (*65*). Freeze-killed 3- to 6-d-old wild-type virgin females were used as tester females. A couple of one male and one female was transferred into one well of an observation chamber (15 mm diameter, 3 mm depth). Then, courtship behaviors were videotaped for 10 min. The courtship index (CI) was calculated as: CI (%) = courting time (s) / 600 (s) × 100. The performance index (PI) was calculated using the following equation to evaluate the reduction rate of CI: PI = (average CI_control_ – average CI_stress_) / average CI_control_.

In measuring spontaneous locomotor activity, a male was placed into one well of an observation chamber. Spontaneous locomotion was videotaped for more than 10 min. To reduce variability, the background was subtracted using a customized script of MATLAB (2019a, MathWorks, USA). The total travel distance from 5 to 605 s was measured using tracking software (Move-Tr/2D, Library, Japan).

In measuring feeding behavior, 20 to 30 virgin males were kept in separate vials until the experiment. For starvation induction, males were transferred to vials with 1% agarose gel from 23 to 25 h before the feeding assay. After SS stress, the 1 h feeding behavior of males was monitored using FlyPAD (*32*). Electrodes on both sides in each arena were filled with 1% agarose gel with 10% sucrose as a food source. As indexes of the appetite of flies, the number of sips and sip duration were used in this study. The numbers of sips detected in two electrodes were summed to obtain the total number of sips. For sip duration, the measured sip durations in two electrodes were averaged.

### 3IY feeding

For the inhibition of *TH* at an adult stage, CS males were fed with 0.1 or 1.0 mg/ml 3IY (I0075, Tokyo Chemical Industry, Japan) from 2 d before the experiments. After 1 h SS stress, control and 3IY-fed males were kept in breeding vials containing food except for the case of immediate tests.

### Temporal disruption of neurotransmission by Shi^ts1^

*shi^ts1^* was driven by four GAL4 lines (*TH*-GAL4, *R58E02*, *MB504B*, and *NP5945*). In this study, the RT and PT were 30 °C and 25 °C, respectively, except in the experiment using *TH*-GAL4. In the experiment using *TH*-GAL4, genetic controls and F_1_ hybrids were reared and used in experiments at 20 °C to minimize the effects of Shi^ts1^ at PT. The temperature of all chambers was preadjusted to RT and PT. All courtship measurements were conducted at 25 °C.

### Temporal activation of dopamine neurons by CsChrimson

*CsChrimson* was driven by three GAL4 lines (*R58E02*, *MB504B,* and *TH*-GAL4). For the simultaneous activation of PPL1 and PAM, the F_1_ hybrid of *MB504B* and UAS-*CsChrimson*, *R58E02*, was used. At least 2 d before the experiments, each male was transferred into a small rearing vial with 400 µM ATR-containing food or regular fly food. Males transferred into vials with regular fly food were used as controls. ATR-containing food was prepared by dissolving ethanol with 100 mM ATR (R2500, Sigma-Aldrich, USA) into regular fly food. The small rearing vials were placed in a black box until the experiments. All procedures were carried out in semidarkness (< 2.0 μW/cm^2^). Red light (0.5–1.0 mW 630 nm LED, WSE-5510 VariRays I, Atto, Japan) was irradiated onto small rearing vials to activate dopamine neurons. In each experiment using red light, the irradiance at the center of the irradiated range was measured using an irradiance meter (HD2302.0, Delta Ohm, Japan) and adjusted to 1000 ± 50 µW/cm^2^ with a voltage regulator. After 1 h red light irradiation, males were kept in a black box for 1 h. Subsequently, male courtship activity was measured in semidarkness. PI was calculated using the average CIs of ATR-fed and non-ATR-fed males.

### Optogenetic activation in MB γ neurons after SS stress

*CsChrimson* was driven in MB γ neurons by *R72B08*-GAL4. Males were transferred into small rearing vials with 400 µM ATR-containing food or regular fly food at least 2 d before the experiments. The small rearing vials were placed in a black box until the experiments. All procedures were carried out in semidarkness (< 2.0 μW/cm^2^). For SS stress, one male was transferred into one well of a standard chamber or an SS chamber in a black box for 1 h. Subsequently, males were returned to small rearing vials and kept there with or without 10.0 ± 0.5 µW/cm^2^ red light. Then, male courtship activity was measured in semidarkness. PI was calculated using the average CIs of control and stressed males.

### Immunohistochemistry

The fly brains were dissected in ice-chilled PBS. The dissected brain samples were fixed with 4% formaldehyde for 20 min at room temperature. The fixed brains were washed three times for 20 min with 0.2% Triton-X 100 and incubated with 1% normal goat serum (NGS) for 1 h. Primary antibody staining was conducted using 1:200 mouse anti-TH antibody (22941, ImmunoStar, USA) for 2 d at 4 °C. Then, the samples were washed thrice for 20 min with 0.2% Triton-X 100 and blocked for 30 min with 1% NGS. Secondary antibody staining was conducted with 1:1000 anti-mouse IgG conjugated to Alexa Fluor 568 (A-11004, Thermo Fisher Scientific, USA) for 2 d. After washing three times for 20 min with 0.2% Triton-X 100, the brain samples were mounted with phosphate-buffered saline (PBS) and observed under a confocal microscope (C2, Nikon, Japan). Fluorescence was excited by a 561 nm laser, and images were obtained with a 20× objective lens (Plan Apo VC 20× DIC N2, Nikon, Japan). The pinhole size was 20 µm, and the z-interval was 0.85 µm. Samples from all genetic controls and F_1_ hybrids between *nSyb*-GAL4 and UAS-*TH* RNAi lines were simultaneously processed using the same solutions.

### CaMPARI2 imaging

*CaMPARI2* was expressed by five GAL4 lines (*TH*-GAL4, *R58E02*, *NP6024*, *R35B12*, and *R72B08*). For the Ca^2+^ imaging during SS stress, naive (control) and stressed males were irradiated with UV light for 1 h. For the Ca^2+^ imaging for 10–70 min after the SS stress experience, males were transferred to food-containing chambers after 1 h SS stress. For Ca^2+^ imaging for 70–130 min after the SS stress experience, males were kept in small rearing vials for 1 h after the stress experience. Subsequently, males were transferred to food-containing chambers. 395 nm LED (B09YDJC5DX, Amazon, USA) was used as a UV light source. In each experiment, UV light was irradiated for 1 h. UV light irradiance was measured by an irradiance meter (HD2302.0, Delta Ohm, Japan). Irradiances at the center of the irradiated range were as follows: 20.0 ± 0.2 mW/cm^2^ for PAM, 15.0 ± 0.2 mW/cm^2^ for the other clusters of dopamine neurons, and 10.0 ± 0.2 mW/cm^2^ for MB lobes. After UV light irradiation, males were immediately ice-anesthetized, and then dissections were conducted in 5 mM ethylene glycol-bis(β-aminoethyl)-N,N,Nʹ,Nʹ-tetraacetic acid (EGTA)-containing PBS within 2 h. The extracted brains were immediately mounted with 5 mM EGTA-containing PBS and observed under a confocal microscope (AX, Nikon, Japan). CaMPARI2 was excited by 488 nm and 561 nm lasers, and images were obtained with a 20× objective lens (Plan Apo VC 20× DIC N2, Nikon, Japan). To reduce the image-capturing time and minimize photobleaching, the pinhole size and z-interval were set to large values. In the imaging of dopamine neurons, the pinhole size was 39.11 µm and the z-interval was 1.218 µm. In the imaging of MB neurons, the pinhole size was 153.26 µm and the z-interval was 13.212 µm. For the measurement of the photoconversion rate of CaMPARI2, ROIs were set according to the following method. For dopamine neurons, a round ROI was set on each soma along a particular z-plane using a customized script of MATLAB (2019a, MathWorks, USA). The x–y–z position and radius of ROIs were manually determined to roughly maximize the total fluorescence intensity and minimize the background and the overlapping with other neurons. For MB lobes, an acquired 3D image was projected onto a single plane and analyzed using NIS-Elements AR 4.50.00 (Nikon, Japan). A polygonal ROI was set to surround each lobe. The total red and green fluorescence intensities within each ROI were summed, and the red/green fluorescence ratio was used as an index.

### Real-time quantitative reverse transcription PCR (qRT-PCR)

qRT-PCR analyses were carried out as previously described with some modifications (*66*). Total RNA was extracted from 30–40 heads of 3–8-d-old males using TRIzol (15596026, Thermo Fisher Scientific, USA) as one sample (210–320 heads were used for each genotype). cDNA was synthesized using an RT reagent kit with gDNA Eraser (RR047A, Takara Bio, Japan). qRT-PCR was carried out using THUNDERBIRD SYBR qPCR Mix (QPS-201, TOYOBO, Japan) and LightCycler 96 (05815916001, Roche, Swiss). The mRNA level of a target gene in each sample was measured as a ratio to the *rp49* mRNA level for the internal control. Then, the ratio of the mRNA levels of the target gene to that of *rp49* in each genotype was divided by that in *nSyb*-GAL4/+ flies for normalization. The following primers were used in the experiments: *Dop1R1*-Forward, 5′-TAGCGATTGCGGATCTCTTCG-3′; *Dop1R1*-Reverse, 5′-TGACATCAAAGGCCACCCAAG-3′; *Dop1R2*-Forward, 5′- TCGATAGAGAGAGCGAGTAGAGG-3′; *Dop1R2*-Reverse, 5′- TGATTCTGTTCCTGTTCCAATTTCC-3′; *Dop2R*-Forward, 5′-TCGCTGAGCAGCTTCTACATAC-3′; *Dop2R*-Reverse, 5′-CGTGAGTTCCGATAGGTGGG-3′; *rp49*-Forward, 5′- AAGATCGTGAAGAAGCGCAC-3′; *rp49-*Reverse, 5′-TGTGCACCAGGAACTTCTTG-3′.

### Statistical analyses

All the statistical analyses were performed using IBM SPSS Statistics (version 26 and 28, IBM, USA) except for the analyses of PI. All comparisons were performed as a two-sided test. In all statistical analyses except for PI, the Kolmogorov–Smirnov test and Levene’s test were used to test the normality and homoscedasticity, respectively. Non-normal distributions were mechanically transformed according to logarithmic, exponential, and square root transformations, and then normality was retested. In the statistical analysis of CI, all distributions did not show coherent normality even after the transformations. Thus, all data were processed using the nonparametric Mann–Whitney *U* test. A randomization test based on the bootstrapping method was used in the statistical analysis of PI with an open R script (*67*). Briefly, random resampling of CI was executed within each group. PI was calculated using bootstrapped control CI and SS stress CI, and then PI in each group was compared with that of the control group. After comparing 10,000 times, the *P* value was determined as a fraction of lower or higher PI than that of the control. The *P* value was doubled owing to the two-sided test.

In the statistical analysis of spontaneous locomotor activity, the Mann–Whitney *U*-test was used for travel distances immediately after 1 h SS stress and 1 d after 24 h SS stress and Student’s *t*-test following to logarithmic transformation was used for travel distances 1 h after 1 h SS stress and 1 d after 7 h SS stress. In the statistical analysis of feeding activity, the number of sips and sip duration were processed by the Student’s *t*-test as distributions showing normality and homoscedasticity. In the statistical analysis of Ca^2+^ imaging of dopamine neurons during 1 h SS stress, the Mann–Whitney *U*-test was used for PAM, PAL, PPL1, PPL2c, PPM2, and PPM3, and Student’s *t*-test was used for PPL2ab with square root transformation, PPM1 and T1. In the statistical analysis of Ca^2+^ imaging of dopamine neurons after 1 h SS stress, the Mann–Whitney *U*-test was used for PPL1 and PPL2c, Welch’s *t*-test for PAM with logarithmic transformation and PPL2ab with square root transformation, and Student’s *t*-test was used for PPM1. In the statistical analysis of Ca^2+^ imaging of MB lobes during 1 h SS stress using non-3IY-fed males, Welch’s *t*-test was used for the β lobe with logarithmic transformation, and Student’s *t*-test was used for the γ lobe, the α lobe with logarithmic transformation, and the α’ and β’ lobes. In the statistical analysis of Ca^2+^ imaging of MB lobes during 1 h SS stress using 3IY-fed males, Welch’s *t*-test was used for the α lobe, the β lobe, and the α’ lobe with exponential transformation, and Student’s *t*-test was used for the γ and β’ lobes. In the statistical analysis of Ca^2+^ imaging of MB lobes for 10–70 min after 1 h SS stress, the Welch’s *t*-test was used for the γ lobe of 3IY-fed males and Student’s *t*-test was used for the γ lobe, the α lobe, the β lobe, the α’ lobe with exponential transformation, and the β’ lobe with square root transformation for the non-3IY-fed males. In the statistical analysis of Ca^2+^ imaging of the MB γ lobe for 70–130 min after 1 h SS stress, the Student’s *t*-test was used. In the statistical analysis of qRT-PCR results, all distributions showed normality. In *Dop1R1* and *Dop1R2* KD, distributions did not show coherent homoscedasticity. Thus, data were processed using the Games–Howell test following Welch’s ANOVA. In *Dop2R* KD, distributions showed homoscedasticity. Thus, data were processed by the Tukey HSD test following one-way ANOVA.

## Acknowledgments

We thank Kahori Sasaki and Emiko Nakagawa for their technical assistance and Kohei Ueno for carefully reading the manuscript and providing critical comments. We also thank Shoma Sato and Show Inami for their helpful discussions. We are grateful to the Bloomington Drosophila Stock Center and the Drosophila Genomics and Genetic Resources (DGGR, Kyoto Stock Center) for providing the fly strains.

## Author contributions

Conceptualization: TSato, TK, TSakai

Methodology: TSato, TK, TSakai

Investigation: TSato, RT

Visualization: TSato

Supervision: TSakai

Writing—original draft: TSato, TK, TSakai

Writing—review & editing: TSato, TK, Tsakai

## Data Availability

All relevant data is within the paper and its supplementary materials.

