## Supplementary material for "Confinement stress with movement restriction suppresses male courtship in *Drosophila* through dopamine-dependent neuroplasticity": Supplimental Figures

### Supplementary Figures

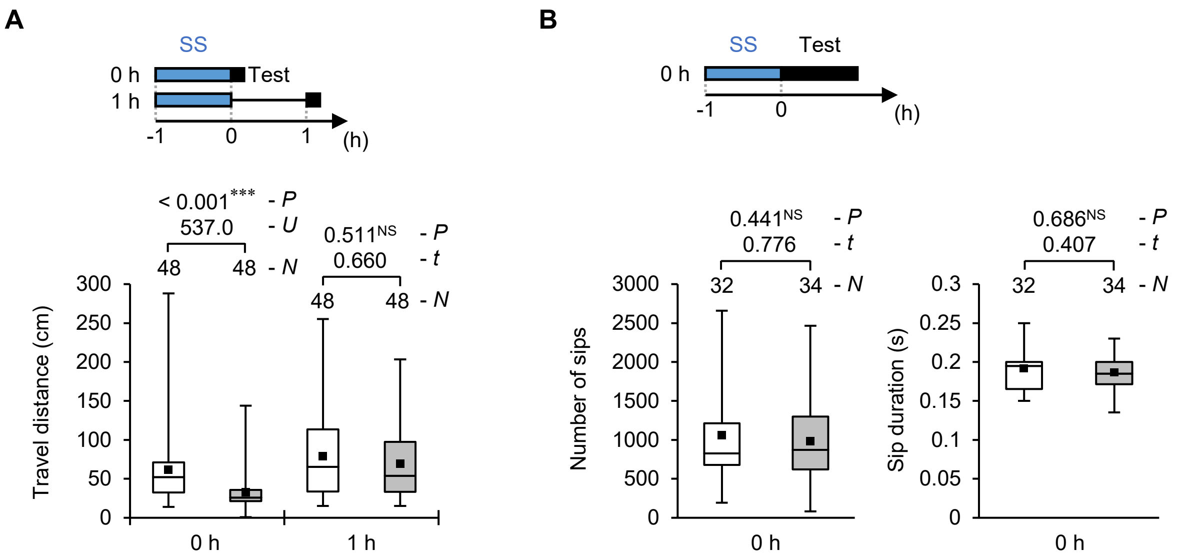


**Fig. S1 Spontaneous locomotor activity and feeding behavior are intact 1 h after 1 h SS stress.** (**A**) To measure general locomotor activity, the total travel distances for 10 min were measured immediately and 1 h after 1 h SS stress. (**B**) Feeding behavior was observed for 1 h after 1 h SS stress. The number of sips and sip duration in each male was measured using FlyPAD. (**A** and **B**) Wild-type (CS) males were used in the experiments. In each graph, white boxes indicate naive males and gray boxes indicate stressed males. Box plots show 0th, 25th, 75th, and 100th centiles. In the box plots, black squares indicate the mean, and the lines are drawn at the median. For statistical comparisons, the Mann–Whitney *U*-test was used for the travel distance immediately after 1 h SS stress. Student's *t*-test was used for the travel distances 1 h after 1 h SS stress and the number of sips and sip duration. ***, $P < 0.001$; NS, not significant.

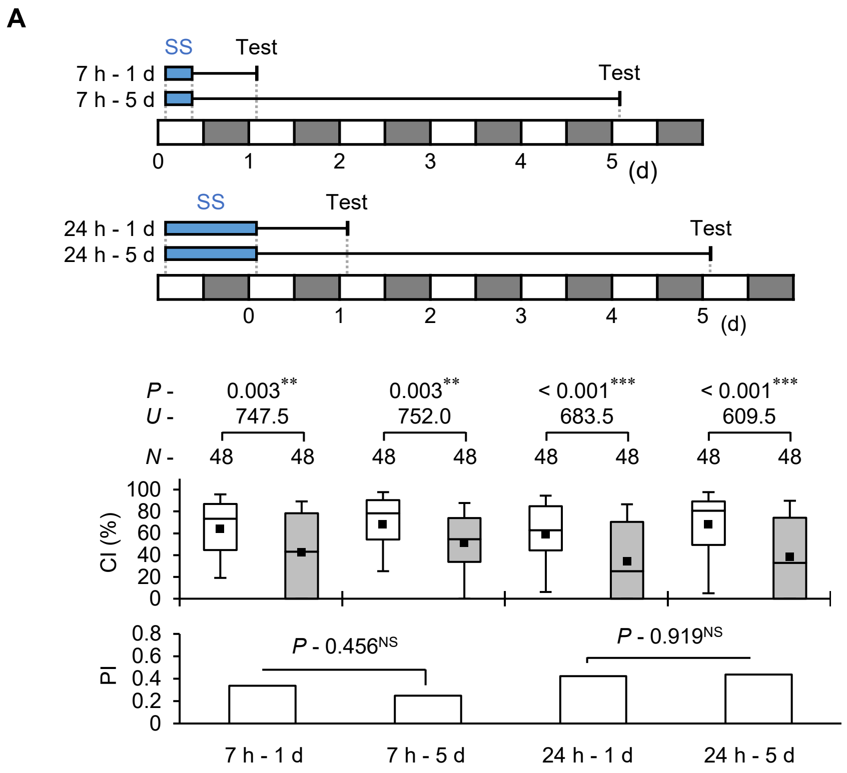


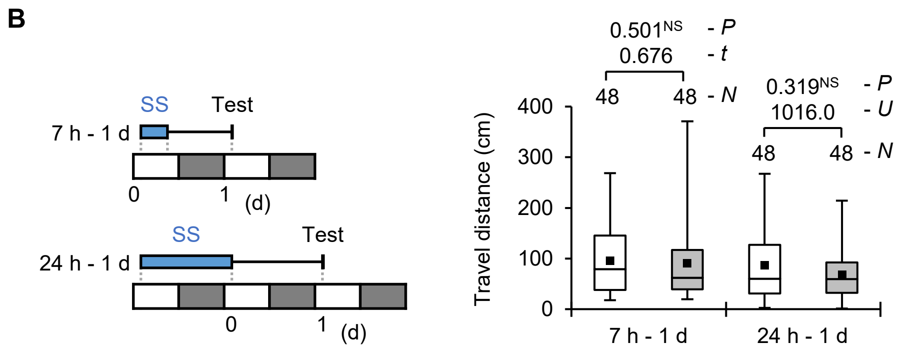


**Fig. S2 7 h and 24 h SS-stress-induced courtship suppression lasts at least 5 d.** (**A**) Courtship activity was measured 1 or 5 d after 7 or 24 h SS stress. (**B**) Total travel distances during 10 min spontaneous locomotor were measured 1 d after 7 or 24 h SS stress. (**A** and **B**) Wild-type (CS) males were used in the experiments. In each graph, white boxes indicate naive males and gray boxes indicate stressed males. Box plots for a set of CI data show the 10th, 25th, 75th, and 90th centiles, and those for a set of travel distance data show the 0th, 25th, 50th, 75th, and 100th centiles. In the box plots, black squares indicate the mean, and the lines are drawn at the median. For statistical comparisons, the Mann–Whitney *U*-test was used for CI and travel distance 1 d after 7 h SS stress. Student's *t*-test was used for travel distance 1 d after 7 h SS stress. A bootstrapping-based randomization test with an R-script was used for PI. ***, $P < 0.001$; **, $P < 0.01$; NS, not significant.

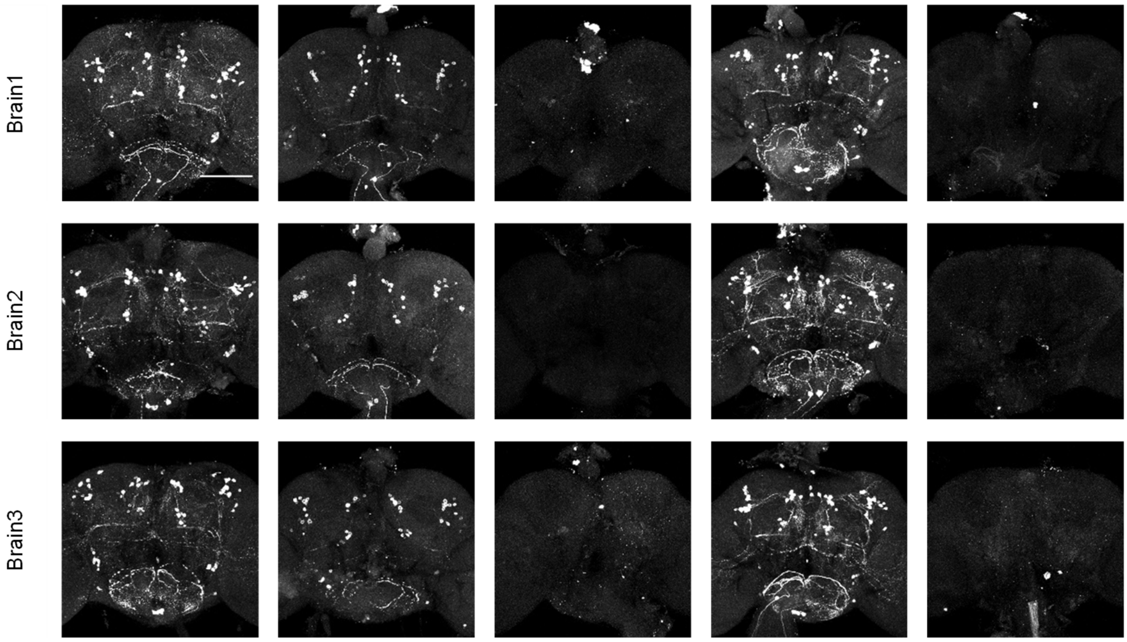


| | | | | | |
|---|---|---|---|---|---|
| *nSyb*-GAL4 | + | - | + | - | + |
| UAS-*TH* RNAi #1 | - | + | + | - | - |
| UAS-*TH* RNAi #2 | - | - | - | + | + |

**Fig. S3 Antibody staining of the adult brains in TH knockdown males.** Representative images of male brains stained with an anti-TH antibody are shown for each genotype. All confocal images of the adult brain were taken from the posterior side. Scale bars indicate 100 μm.

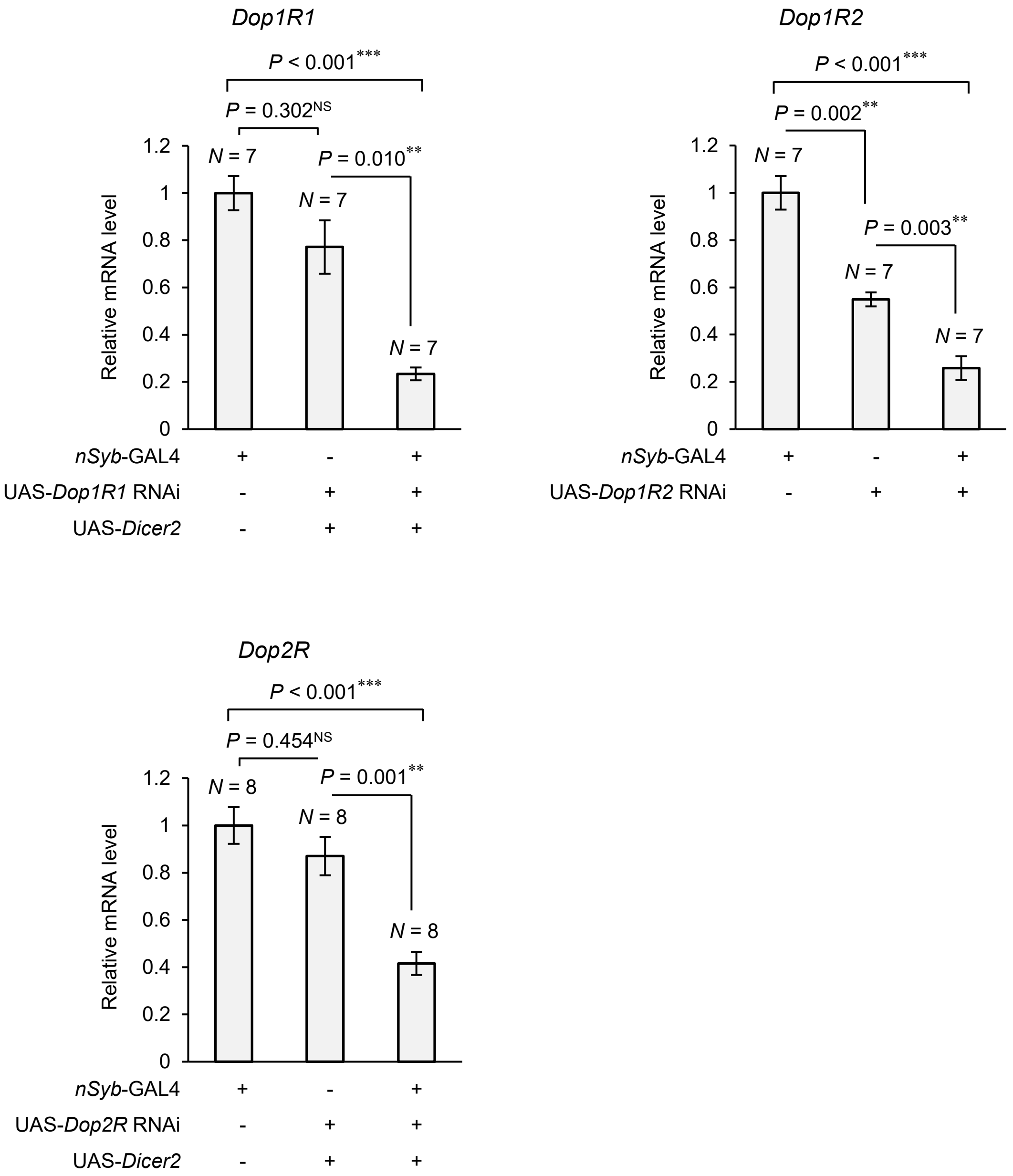


**Fig. S4 Real-time qRT-PCR analysis of *Dop1R1*, *Dop1R2,* and *Dop2R* knockdown**. The relative mRNA expression levels of *Dop1R1*, *Dop1R2*, and *Dop2R* in each genotype were measured. Total RNA was extracted from 30–40 male heads as one sample (210–320 heads were used for each group). Bar plots show the average ± SEM of each distribution. For statistical comparisons, the Games–Howell test following Welch's ANOVA was used for the comparison of relative *Dop1R1* and *Dop1R2* mRNA expression levels, and the Tukey HSD test following one-way ANOVA was used for the comparison of relative *Dop2R* mRNA expression levels. ***, $P < 0.001$; **, $P < 0.01$; NS, not significant.

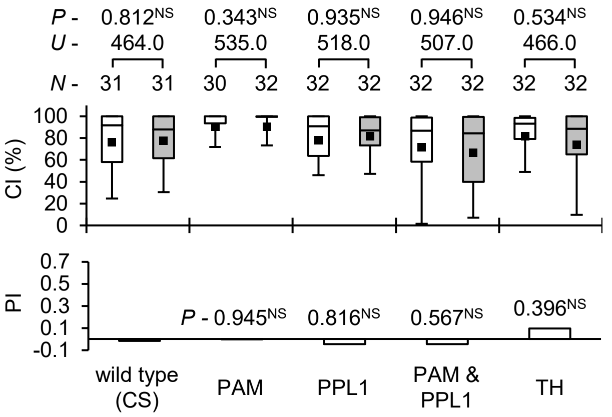


| | wild type (CS) | PAM | PPL1 | PAM & PPL1 | TH |
|---|---|---|---|---|---|
| *R58E02*-GAL4 | - | + | - | + | - |
| *MB504B*-GAL4 | - | - | + | + | - |
| *TH*-GAL4 | - | - | - | - | + |
| UAS-*CsChrimson* | - | + | + | + | + |

**Fig. S5 ATR feeding does not affect male courtship activity**. Flies were maintained in semidarkness for more than 2 d before the start of the experiments; and subsequently, their courtship activity was measured for 10 min in semidarkness. Wild-type (CS) males and males expressing CsChrimson in a subset of dopamine neurons were used in the experiments. Three GAL4 lines (*R58E02* for PAM, *MB504B* for PPL1, and *TH*-GAL4 for almost dopamine neurons) were used. In each graph, white boxes indicate naive males and gray boxes indicate ATR-fed males. Box plots for a set of CI data show the 10th, 25th, 75th, and 90th centiles. In the box plots, black squares indicate the mean, and the lines are drawn at the median. For statistical comparisons, the Mann–Whitney *U*-test was used for CI. For PI, bootstrapping-based randomization test with an R-script was used. *N*, sample size; *** $P < 0.001$; NS, not significant.

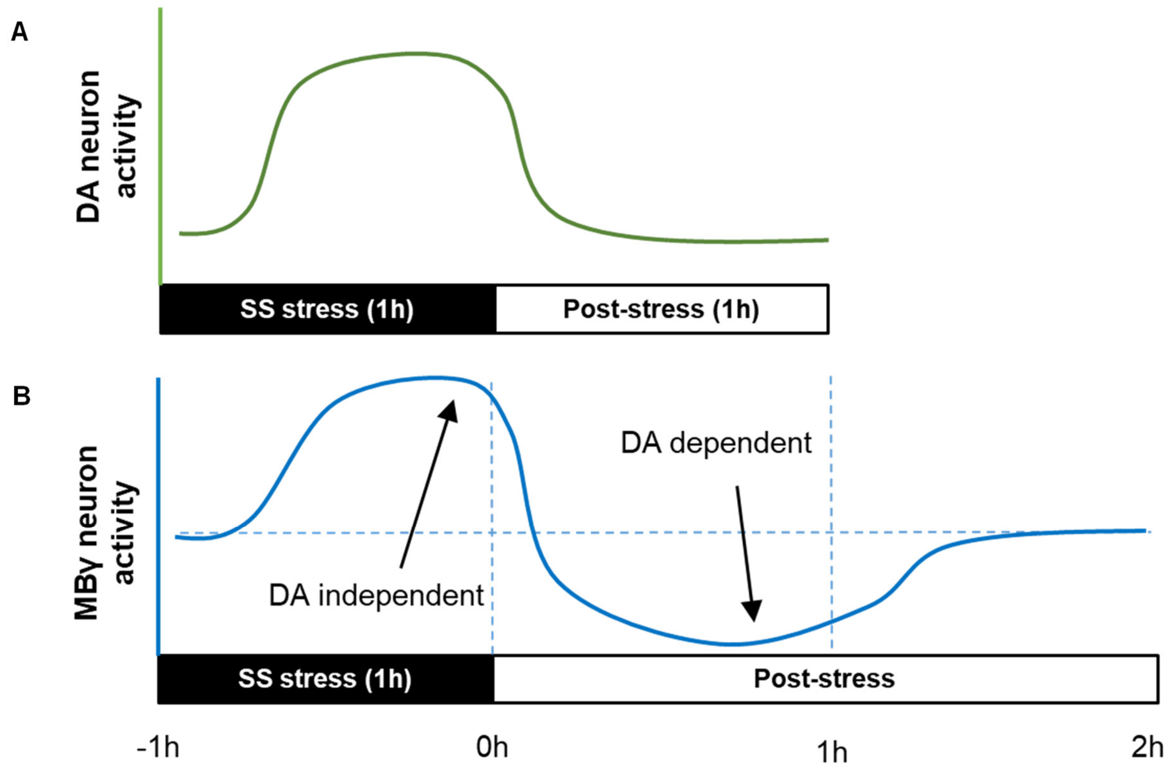


**Fig. S6 A possible model of fluctuating activity of dopamine neurons and MB during and after SS stress experience**. (**A**) Schematic of fluctuating activity of dopamine neurons in a stress-dependent manner. (**B**) Schematic of fluctuating activity of MB neurons during and after stress experience.
